# Tunable light-focusing behavior of engineered bacterial microlenses with controllable shapes

**DOI:** 10.64898/2025.12.17.694918

**Authors:** Lynn M. Sidor, Kathren P. Sage, Michelle M. Beaulieu, Dylan F. Wilson, Emerson Jenen, B. Cansu Acarturk, Wil V. Srubar, Greg R. Schmidt, Elio A. Abbondanzieri, Anne S. Meyer

**Affiliations:** Department of Biology, University of Rochester, Rochester, New York, 14627, USA; Department of Physics and Astronomy, University of Rochester, Rochester, New York, 14627, USA; Department of Biochemistry and Biophysics, University of Rochester Medical Center, Rochester, New York, 14627, USA; Dept. of Civil, Environmental, and Architectural Engineering, University of Colorado Boulder, Boulder, Colorado, 80309, USA; Materials Science and Engineering Program, University of Colorado Boulder, Boulder, Colorado, 80309, USA; Institute of Optics, University of Rochester, Rochester, New York, 14627, USA

## Abstract

Recently, engineered bacterial cells have been shown to behave as optically-active photonic devices comparable to industrially fabricated microlenses^1^. Bacterial cells can be encapsulated within a layer of polysilicate through surface display of the sea sponge enzyme silicatein, which mineralizes a polysilicate coating. The addition of this polysilicate layer significantly enhances the ability of these cells to guide, scatter, and focus light^1^. However, this previous technique was limited to creating rod-shaped microlenses, which are not ideal for all applications. Here we expand upon this technology by engineering the shapes of silicatein-displaying bacterial cells. Through the overexpression of the genes *bolA*^2–5^ and *sulA*^6,7^ or through the use of the drug A22^8,9^, we are able to alter *Escherichia coli* cells from their characteristic rod-like shape to either spherical or filamentous forms. Round cells encapsulated in polysilicate were shown to scatter light more intensely and symmetrically than rod-shaped cells, while encapsulated filamentous cells were shown to guide light similarly to an optical fiber. This control over the size and shape of optically-active cells is a major advancement towards developing bio-engineered photonic devices such as nanophotonic waveguides, spherical microlens arrays, and advanced biosensors.

## Introduction

Nanophotonic devices, which can manipulate light over sub-wavelength volumes, show great potential in the development of a broad range of biosensors capable of overcoming the limitations of sensitivity, throughput, ease-of-use, and miniaturization inherent to current bio-analytical methods^10^. Recently, utilizing a synthetic biology approach, we have demonstrated that bacterial cells can be engineered to act as microlenses^1^. These *Escherichia coli* bacterial cells were genetically modified to surface-display a single enzyme derived from sea sponges, silicatein, via fusion to the outer membrane protein OmpA. In the presence of aqueous silica precursor molecules, strains expressing silicatein from *Tethya aurantia* (TaSil) or *Suberites domuncula* (SdSil) were able to polymerize a layer of polysilicate, also known as bioglass, surrounding the bacterial cell. These polysilicate-encapsulated cells behaved like microlenses by focusing light several micrometers away from the edge of the cell, with higher peak intensity than seen for a wild-type *E. coli* cell. These microlenses adopted the rod-shaped form characteristic of *E. coli* cells, *i.e.*, a cylinder with hemispherical ends approximately 1-3 µm long and 0.8 µm in diameter. This shape may be well-suited for some applications requiring directional focusing and a specific size range but is more poorly suited for applications requiring directionally homogenous focusing or applications that require larger cell lengths. Control over the cell shape would make it possible to tailor the optical properties of polysilicate-encapsulated cells. Engineering more spherical cells would be useful for creating lenses that have symmetric focal properties regardless of the direction of incoming light. Conversely, longer cell shapes would be more ideal for creating light guides, similar to a fiber optic, that could transmit light over long distances.

The shape of bacterial cells can be altered via numerous pathways, both genetically and through the use of specific drugs that target key processes within the cells. Previous work has established that *E. coli* cells can be converted from rod-shaped to more spherical shapes through overexpression of the gene *bolA*^2–5^. BolA has been identified as a stress response protein and has been linked to biofilm formation and the regulation of several cell elongation and division proteins, including MreB^4^. MreB is an actin homolog involved in maintenance of the rod shape of *E. coli* cells. When the spatial organization of MreB filaments is disrupted through BolA overexpression, the cell cannot localize the cytoskeletal elements necessary to create the typical rod shape of the cell, resulting in a cell with a more spherical shape^4^. The drug S-(3,4-dichlorobenzyl)isothiourea, also referred to as A22^8,9^, has also been shown to perturb MreB activity. A22 can directly bind to MreB as a competitive inhibitor of ATP, which reduces the ability of MreB to polymerize^8^ and creates spherical *E. coli* cells.

To produce elongated cells, *E. coli* cells can be induced to grow filamentously, in which the cells grow much longer than the typical ~1.5-3 µm cell length. Filamentous growth can be accomplished by either genetic manipulation or treatment with a drug, including via *sulA* overexpression^6,7^ or cephalexin treatment^11–14^ respectively. SulA is an SOS response protein that directly interacts with the cell-division protein FtsZ, blocking the assembly of Z-rings and thereby causing the *E. coli* to undergo filamentous growth^6,7,13^. Cephalexin is a FtsI-specific β-lactam antibiotic that inhibits FtsI, the only essential cell division protein^12–15^, which triggers the cells to grow without undergoing cell division. With these approaches, bacterial cells can be induced to grow into highly elongated, fiber-like shapes.

In this work, we have combined shape manipulation with polysilicate encapsulation of our engineered bacterial cells. Experimental conditions were identified in which the bacterial cells demonstrated altered spherical or elongated shapes while also displaying polysilicate localization to their outer surfaces. These cells showed structural changes to their cell surfaces, as previously reported for polysilicate-encapsulated rod-shaped cells, and they remained metabolically active for several months following the encapsulation and shape alteration treatments. Both the spherical and elongated polysilicate-encapsulated cells were able to scatter and focus light. The spherical cells focused light with higher intensity than the rod-shaped cells, and the scattered light was not dependent on the orientation of the cells due to their nearly spherical shapes. The scattering observed for the elongated cells was brighter than for any other cell shapes measured and was highly cell-orientation dependent, with cells that were most aligned to the incoming light able to transmit light more than 10 µm along the length of the cell body. We have created bacterial cells that have potential applications as tunable photonic devices. These cells have adjustable optical properties that can be used for diverse applications including microlens arrays, photonic nanojets, and microscopic waveguides. The ability to use bacterial cells to self-assemble into spherical or elongated polysilicate-encapsulated nanophotonic devices has the potential to make the production of miniaturized microlenses and nanophotonic light guides both more cost-effective and more biocompatible for integration into medical diagnostic devices.

## Results

### Light focusing by rod-shape cells is cell-orientation dependent

Typical *Escherichia coli* cells are rod-shaped and approximately 1.8 x 0.8 μm in size, such that their length is greater than their width. Therefore, the light-scattering and focusing behaviors of engineered polysilicate-encapsulated *E. coli* cells could potentially be affected by their orientation in relation to the angle of light. To analyze whether cell orientation influences the light scattering of rod-shaped polysilicate-encapsulated cells^1^, we illuminated TaSil and SdSil bacteria at a range of incident angles from −90° to 90° via Multiple Angle Illumination Microscopy (MAIM) and measured the light scattered by the bacteria cells across the range of illumination angles. Maximum intensity projections of the scattered light were analyzed as a function of distance from the bacteria cells, and this data was divided into bins based upon the angle of the long axis of the cell in relation to the incoming light. Bins were delineated as: 0-30°, with cell poles oriented towards the incoming light; 31-60°, with cells oriented diagonally relative to the incoming light; and 61-90°, with the cell poles oriented perpendicularly to the incoming light (**Figure 1A**). Both TaSil and SdSil cells from every bin of orientation were observed to scatter bright jets of light (**Figure 1B-D**), and the scattered light for each bin exhibited a peak in average intensity several μm away from the cell, indicative of light-focusing (**Figure 1E**). Comparison of the scattering profiles indicated that cells in the 0-30° bin scattered light with a focal distance of 2.4 µm, while cells in the 31-60° and 61-90° bins scattered light with focal distances of 3.4 µm and 4.3 µm respectively (**Figure 1E**). The profile of the light scattered by cells in the 61-90° bin exhibited a more gradual decrease in intensity at distances farther away from the focal distance compared to cells in the 0-30° and 31-60° bins. No other parameters of the light scattering behavior, including length and width of the scattered jet of light as well as integrated intensity of scattered light, were significantly different between cells in different orientation bins (**Supplemental Figure 1**). This analysis revealed that rod-shaped polysilicate-encapsulated *E. coli* cells demonstrate different scattering profiles when oriented at different angles relative to the illuminating light.

**Figure 1:**
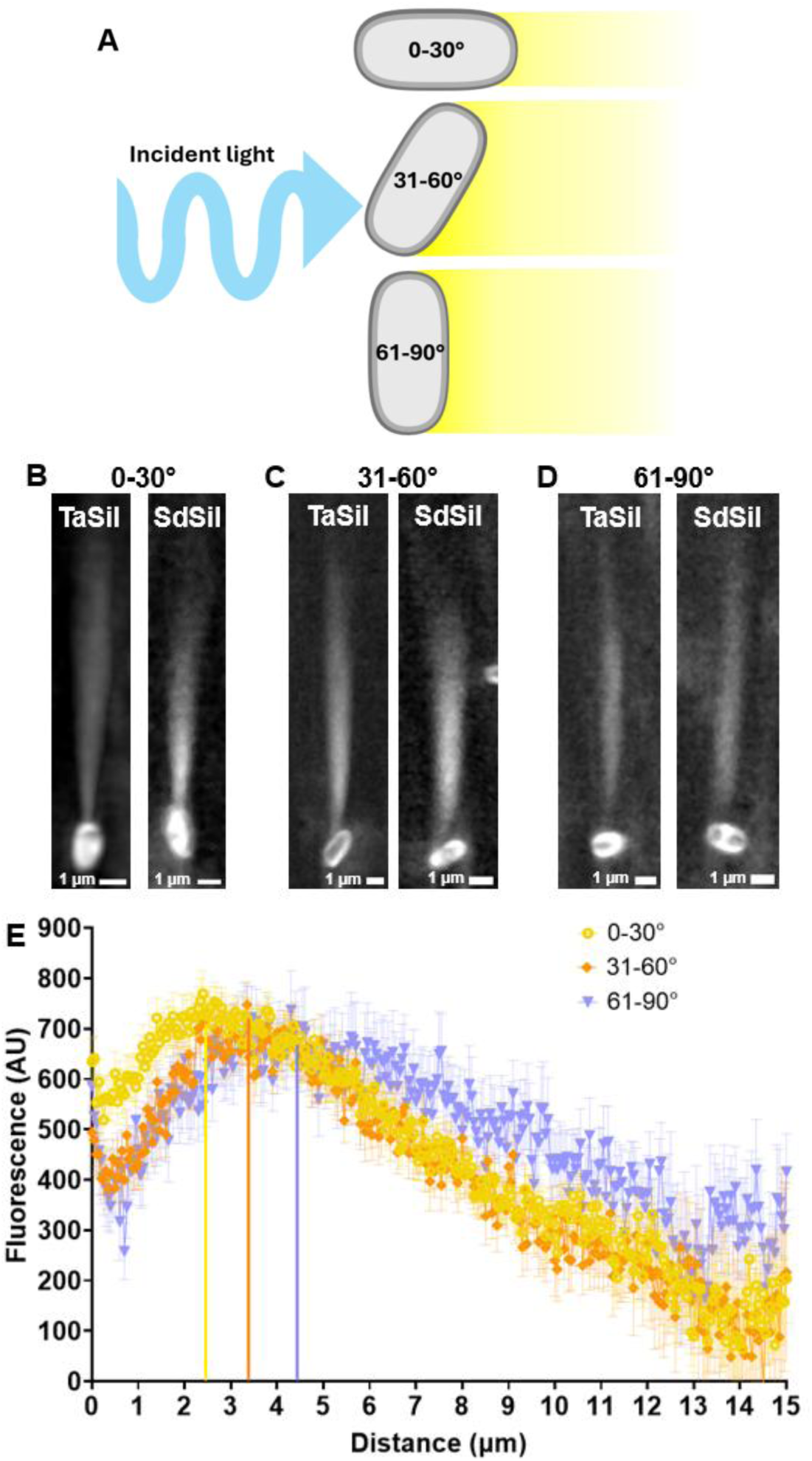
Light scattering by rod-shaped polysilicate-encapsulated bacterial cells is cell-orientation dependent. (A) Schematic illustrating the bins for cell angle in relation to the incident light. (B-D) Maximum intensity projections of individual TaSil and SdSil polysilicate-encapsulated cells scattering light via MAIM. Cells are shown from different bins based on the angle of the long axis of the cell in relation to the incoming light, where (B) is 0-30°, (C) is 31-60°, and (D) is 61-90°. (E) Intensity of scattered light as a function of distance from the edge of the cell, calculated from maximum intensity projections via MAIM, for polysilicate-encapsulated TaSil and SdSil *E. coli* cells oriented at different angles in relation to the incoming light. Vertical lines correspond to the focal peak values for each angle bin. Error bars correspond to standard error of the mean. (n_0-30°_=58, n_31-60°_=33, n_61-90°_=16)

### Construction of inducible spherical, silicatein-expressing strains

Since asymmetrically-shaped, polysilicate-encapsulated *E. coli* cells displayed orientation-dependent light scattering properties, we hypothesized that either decreasing or exaggerating the asymmetry of the cells would result in altered optical properties. In order to produce a bacterial microlens with minimal asymmetry, our goal was to create spherical bacteria cells capable of encapsulating themselves in mineralized polysilicate.

In order to genetically engineer a spherical *E. coli* cell, the protein BolA was placed on an inducible plasmid that was co-transformation compatible with the TaSil and SdSil plasmids. Genetic constructs were assembled containing the BolA protein in several different plasmid backbones and under the control of different promoters in order to test for compatibility with silicatein-expression constructs. The inducible BolA pBAD33 construct was selected for use throughout this work since it was the most consistent at producing spherical cell morphologies when co-induced in silicatein-expressing strains. Attempts at constructing a constitutive BolA-expression strain were unsuccessful, as fully assembled plasmids were found to lack the promoter driving BolA expression following transformation into *E. coli* cells. In parallel, we developed a chemical treatment using the drug A22 to alter the morphology of *E. coli*. Inducible expression of BolA from a plasmid and treatment with A22 were each observed to result in *E. coli* cells converting to more spherical shapes (**Supplemental Figure 2, Supplemental Table 1**).

Cells were fixed at regular timepoints following BolA induction to characterize the progression of the shape-alteration over time, and cell morphology was evaluated via light microscopy. BolA-expressing cells were observed to become increasingly spherical over the time course of BolA inductions (**Supplemental Figure 3A**). Quantification of the circularity of the cells indicated that, prior to BolA induction, similar circularity values were measured for both the BolA cells (0.75 α 0.06) and unmodified wild-type cells (0.78 α 0.06) (**Supplemental Figure 3B**). Following BolA induction, a statistically significant increase in circularity to a value of 0.88 α 0.06 at 3 hours was observed, after which the circularity did not vary significantly for the following 3 hours, with circularity of 0.91 α 0.02 after 6 hours. These data indicated that inducing BolA induction for 3 hours before starting polysilicate encapsulation could allow for the creation of spherical cells that are coated in polysilicate.

To determine whether spherical shape alteration of silicatein-expressing cells is compatible with polysilicate encapsulation, TaSil and SdSil strains carrying the BolA plasmid (TaSil+BolA and SdSil+BolA) were grown in liquid culture, and silicatein expression was induced simultaneously with BolA induction. Following three hours of induction, the cultures were incubated with orthosilicate for three hours. Control samples were only induced for BolA expression but were not induced for silicatein expression. Light microscopy imaging revealed that BolA induction resulted in all silicatein-plasmid-carrying strains becoming more spherical over the course of the induction and encapsulation procedure (**Supplemental Figure 3A**). Circularity quantification indicated that both TaSil+BolA and SdSil+BolA strains reached similar, high levels of circularity by the end of the polysilicate encapsulation procedure, both with (0.91 α 0.02 and 0.89 α 0.03, respectively) and without (0.92 α 0.02 and 0.91 + 0.02, respectively) silicatein induction, with the SdSil strain exhibiting a slight delay in the increase of circularity at earlier timepoints (**Supplemental Figure 3C-D**).

The compatibility of A22-induced shape alteration with polysilicate encapsulation was also investigated. Treatment of wild-type *E. coli* cells with A22 (WT+A22) was observed to convert the rod-shaped cells to more spherical shapes over the course of several hours via light microscopy (**Supplemental Figure 4A**), reaching an average circularity of 0.89 α 0.03 by 3 hours of incubation (**Supplemental Figure 4B**). Silicatein expression was induced simultaneously with A22 addition for both TaSil and SdSil cells (TaSil+A22 and SdSil+A22), and after 3 hours of induction, the cells were incubated with orthosilicate for 3 hours. The TaSil+A22 cells displayed high circularity at the end of the polysilicate encapsulation procedure, both with (0.9 α 0.03) and without (0.91 α 0.02) induction of silicatein (**Supplemental Figure 4A, C**). SdSil+A22 cells demonstrated increased circularity following the polysilicate encapsulation procedure when silicatein was not induced (0.89 α 0.02). However, when silicatein was induced, the SdSil+A22 cells did not demonstrate greatly increased circularity by the end of the encapsulation procedure (0.81 α 0.05) (**Supplemental Figure 4A, D**). Thus, this condition was not used further in this work.

### Spherical cells can stably encapsulate themselves with polysilicate

To investigate whether the spherical silicatein-expressing strains were able to encapsulate themselves in a layer of mineralized polysilicate, as previously observed for cells with unmodified rod shapes^1^, the cells were induced for silicatein expression and shape alteration. Following incubation with orthosilicate, the cells were stained with Rhodamine123, a fluorescent dye that has been used to visualize polysilicate localization^16,17^. Cells were imaged using confocal fluorescence microscopy across multiple months of storage to evaluate the staining pattern for the cells over time. A cell that has a layer of polysilicate on its outer surfaces will display a staining phenotype in which the dye signal is more intense at the cell border and less intense in its center, giving a higher border-to-cytoplasm fluorescence ratio^1^.

For the BolA-expressing strains, the SdSil+BolA strain showed spherical shaped cells with a bright fluorescent signal localized at the cell border with little-to-no internal staining (**Figure 2A**), consistent with peripheral polysilicate localization. The BolA-only control strain demonstrated a dimmer, non-localized Rhodamine123 signal (**Figure 2B**). The TaSil+BolA strain displayed bright areas of staining within the cells, perhaps indicating internal aggregation or inclusion bodies (**Supplemental Figure 5A**). For the A22-treated cells, the TaSil+A22 cells appeared spherical with brightly stained borders and dimly stained interiors (**Figure 2C**), and the WT+A22 cells appeared spherical with diffuse staining (**Figure 2D**). Quantification of the ratio of the border-to-cytoplasm fluorescence signal indicated that SdSil+BolA and TaSil+A22 showed significantly higher ratios than their non-silicatein-expressing control conditions BolA and WT+A22 as well as the TaSil+BolA strain, which all had ratios close to 1 (**Figure 2E, Supplemental Figure 5B**). These data indicate that BolA overexpression in SdSil cells, as well as A22 treatment of TaSil cells, resulted in silicatein-expressing *E. coli* cells with altered, spherically-shaped morphology that were able to encapsulate themselves in a layer of polysilicate.

**Figure 2:**
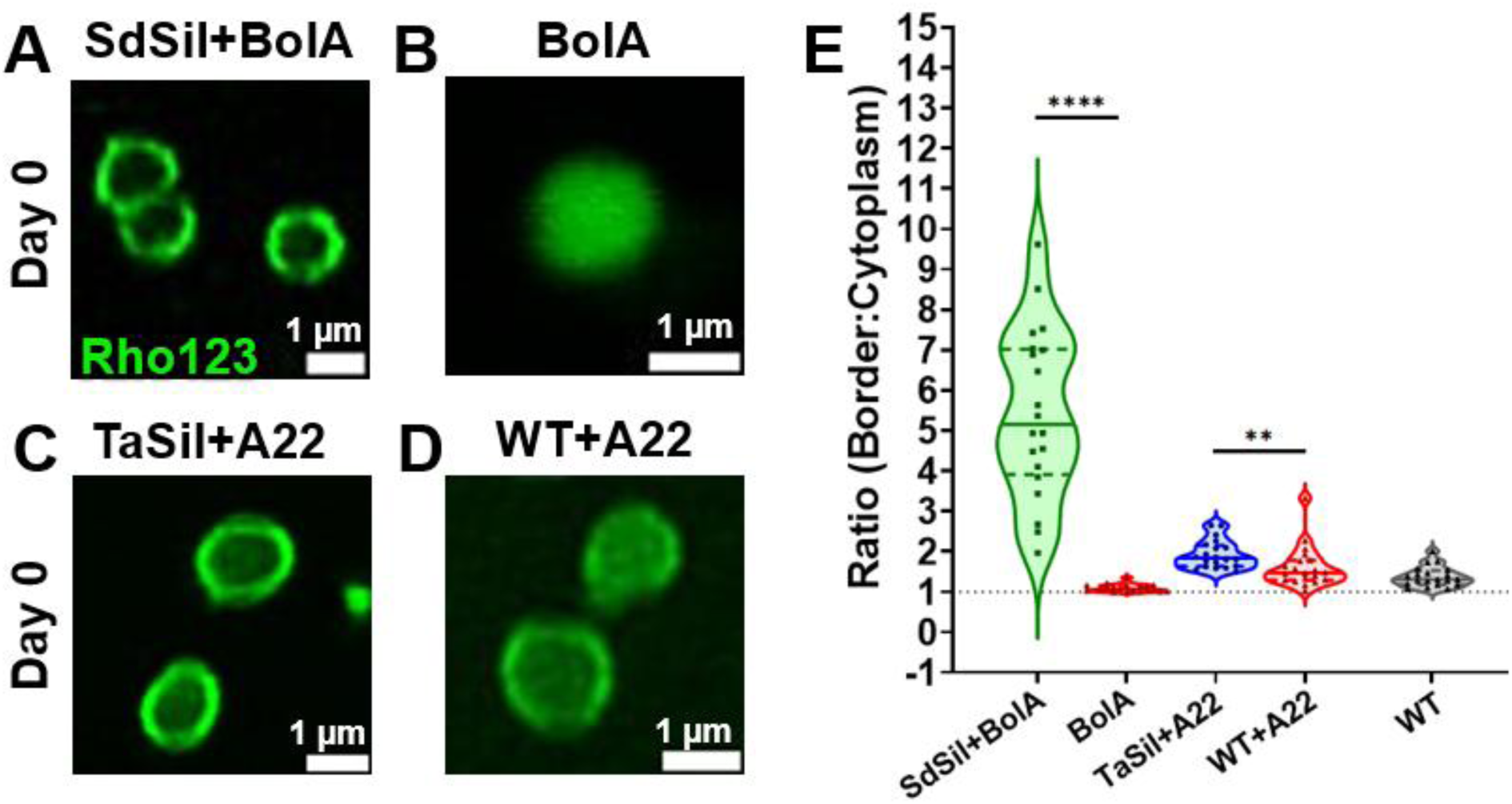
Polysilicate localization on spherical bacterial cells. (A-D) Cells stained with Rhodamine123 at day 0 where (A) are SdSil+BolA, (B) is BolA, (C) is TaSil+A22, and (D) is WT+A22. (E) The quantification of the ratio of Rhodamine123 border to cytoplasm staining for strains A-D and WT. (n=20) ** p≤0.01, **** p≤0.0001.

To assess the polysilicate localization patterns of the spherical cells over time, the samples were incubated in buffer with constant rotation at 4°C, followed by Rhodamine123 staining and confocal fluorescence microscopy at monthly intervals. The SdSil+BolA cells retained a spherical morphology and displayed staining at its outer surfaces at both 1 month (**Supplemental Figure 6A**) and 6 months (**Supplemental Figure 6B**) but showed punctate internal staining at 6 months (**Supplemental Figure 6B**). The ratio of border-to-cytoplasm staining was significantly higher for the SdSil+BolA cells compared to the BolA-only control cells at 1 month (**Supplemental Figure 6A, B, E**). At 6 months, the ratio for SdSil+BolA cells was significantly higher than for wild-type cells but not significantly different from BolA cells, which showed a significantly increased ratio at this timepoint compared to earlier timepoints (**Supplemental Figure 6A, B, E)**. The TaSil+BolA cells maintained a pattern of internal staining and low internal-to-border signal ratio at 1 month and 6 month (**Supplemental Figure 6F and G**). The TaSil+A22 condition showed spherically shaped cells with fluorescent staining at their borders and a significantly higher border-to-cytoplasm ratio in comparison with the WT+A22 control for months 1 and 6, with the intensity of the border staining signal decreasing over time (**Supplemental Figure 6E, H-K**).

To quantify how the altered shapes of the polysilicate-encapsulated spherical cells were maintained over time, the circularity of the cells was evaluated at monthly intervals. The spherical SdSil+BolA and TaSil+BolA strains showed no significant change in their circularity between months 0 and 6, while the BolA-only control strain showed increased circularity over the course of 6 months (**Supplemental Figure 6L**). The TaSil+A22 cells also displayed significantly increased circularity over the 6-month time course, while the WT+A22 control cells did not display significant changes in their circularity (**Supplemental Figure 6L**).

To characterize the morphology of the outer surfaces of the polysilicate-encapsulated spherical cells, thin sections of SdSil+BolA and TaSil+A22 cells were prepared and imaged via transmission electron microscopy (TEM) in comparison to control BolA, WT+A22, and untreated wild-type cells. The polysilicate-encapsulated cells displayed a smoother, less-ruffled cell surface (**Supplemental Figure 7A-B**) than the non-silicatein expressing control cells, which showed ruffled cell surfaces (**Supplemental Figure 7C-E**) typical of unmodified *E. coli* cells^18^. These morphologies were in agreement with the smooth outer borders previously seen for rod-shaped polysilicate-encapsulated strains without shape alterations^1^. Quantification of the surface roughness for the encapsulated strains indicated that their cell surfaces were significantly less rough than for the control, non-encapsulated strains (**Supplemental Figure 7F**). The polysilicate-coated SdSil+BolA and TaSil+A22 cells were also observed to have less electron-dense cell interiors than the control cells (**Supplemental Figure 7G**), again consistent with results from silicatein-expressing cells without shape alteration^1^. Scanning electron microscopy (SEM) of the spherical, polysilicate-encapsulated cells revealed that the SdSil+BolA cells showed a more aggregated, embedded appearance at the high concentrations used for this imaging technique compared to the other strains (**Supplemental Figure 8**).

These data indicate that the SdSil+BolA strain and the TaSil+A22 cells adopted spherical cell shapes and were able to encapsulate themselves in a layer of polysilicate. These cells were able to maintain their spherical shapes and polysilicate coatings for several months. The TaSil+BolA cells did not display a consistent pattern of external polysilicate localization, so this condition was discontinued for the subsequent experiments.

### Spherical, polysilicate encapsulated cells scatter focused light with low-orientation dependence

Since the shape-altered, spherical polysilicate-encapsulated *E. coli* cells displayed highly symmetrical cell shapes, we hypothesized that they would display distinct light-focusing properties. MAIM microscopy revealed that the spherical, polysilicate-encapsulated SdSil+BolA cells scattered a bright jet of light similar in shape to that scattered by the rod-shaped, polysilicate-encapsulated SdSil cells (**Figure 3A**). Interestingly, the BolA control cells, which were spherical but not encapsulated in polysilicate, were also able to scatter a jet of light, whereas the rod-shaped wild-type cells did not display visible scattered light (**Figure 3A**).

**Figure 3:**
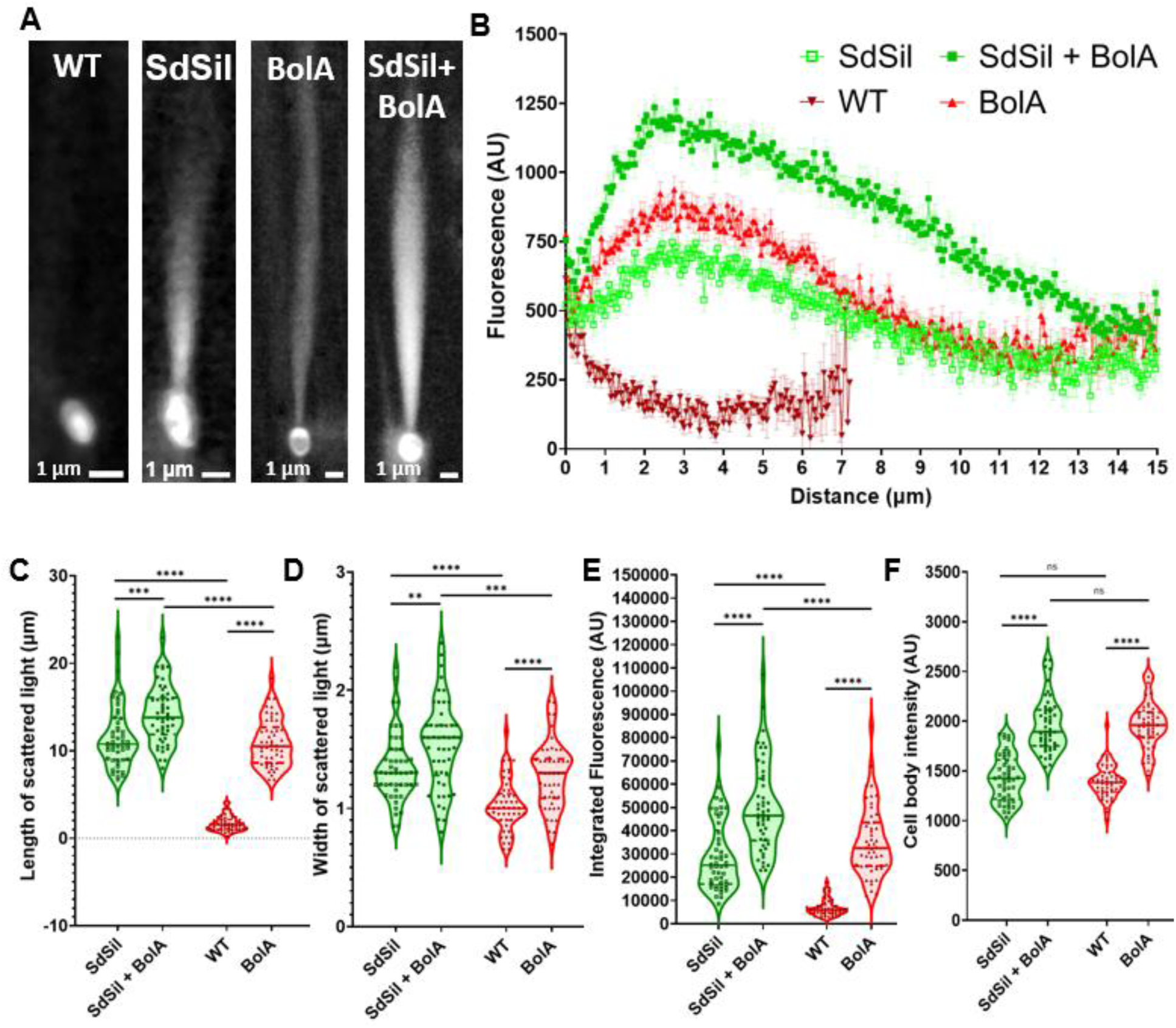
Spherical, polysilicate-encapsulated cells scatter higher-intensity, focused light. (A) Maximum intensity projection for rod-shaped (WT), rod-shaped polysilicate-encapsulated (SdSil), spherical (BolA), and spherical, polysilicate-encapsulated (SdSil+BolA) cells scattering light via MAIM. (B) Profiles of the scattered light as a function of the distanced from the edge of the cell, calculated from the maximum intensity projections, where error bars correspond to standard error of the mean. (C) Length of the scattered light, (D) width of the scattered light, (E) integrated intensity of the scattered light, and (F) mean cell body intensity of the light within cell boundaries, all calculated from the maximum intensity projections. (n=50). ns: no significance, ** p≤0.01, *** p≤0.001, **** p≤0.0001.

Quantification of the scattered light indicated that the SdSil+BolA cells scattered light that was more intense than the rod-shaped SdSil cells throughout the scattering range and decreased in intensity more slowly with increasing distance from the cells (**Figure 3B**). The SdSil+BolA cells displayed a focal peak where the intensity of scattered light was the highest, approximately 2.8 µm away from the cells, which was slightly closer than the focal peak of the SdSil cells. The BolA cells exhibited a scattering profile that was comparable to the SdSil cells, with a focal peak at a similar distance away from the cells and at a slightly higher intensity. The wild-type cells showed lower intensity of scattered light than any of the spherical or polysilicate-encapsulated cells, with no focal peak. Both SdSil+BolA and BolA cells scattered jets of light that were significantly longer (**Figure 3C**) and wider (**Figure 3D**) than their rod-shaped counterparts, with SdSil+BolA cells scattering longer and wider jets than the BolA cells. The integrated intensity of the scattered light was significantly higher for cells from both BolA-expressing strains than for the rod-shaped cells, and SdSil+BolA scattered more light than the BolA cells (**Figure 3E**). The brightness of the spherical SdSil+BolA and BolA cell bodies were not significantly different from each other but were significantly higher than for the SdSil and wild-type cells (**Figure 3F**).

The spherical A22-treated cells were also observed to scatter jets of light, with the polysilicate-encapsulated TaSil+A22 cells scattering brighter jets than the WT+A22 cells (**Supplemental Figure 9A**). The light scattered by the TaSil+A22 cells was lower in intensity than for the rod-shaped TaSil cells across most of the length of the scattered jets of light, with a similar focal distance (**Supplemental Figure 9B**). Similar to the WT+BolA cells, the WT+A22 cells also scattered light that was more intense than the wild-type cells across a distance of several micrometers away from the cells, though less intense than for the TaSil+A22 cells throughout the entire lengths of the jets (**Supplemental Figure 9B**). The jets of light scattered by the TaSil+A22 cells were significantly longer than for the rod-shaped TaSil cells (**Supplemental Figure 9C**), and no significant differences were seen in the width of the jets (**Supplemental Figure 9D**), the integrated intensities of the scattered light (**Supplemental Figure 9E**), or the brightness of the cells (**Supplemental Figure 9F**). The non-polysilicate-encapsulated WT+A22 cells exhibited jets of light that were significantly longer (**Supplemental Figure 9C**) but not wider (**Supplemental Figure 9D**) than for the wild-type cells, with significantly higher integrated intensity of scattered light (**Supplemental Figure 9E**) and higher cell body brightness (**Supplemental Figure 9F**).

Overall, the data show that the induced shape alterations alone resulted in enhanced light-scattering and -focusing properties for the spherical cells, which was further enhanced by polysilicate encapsulation. Both of the spherical, silicatein-expressing strains were able to scatter focused jets of light, with the drug-induced TaSil+A22 performing similarly to the rod-shaped silicatein-expressing cells, and the genetically inducible SdSil+BolA significantly out-performing their rod-shaped counterparts across all light-scattering parameters. The spherical cells displayed significantly lower aspect ratios (**Supplemental Figure 10**), indicating that their light scattering properties have little dependence on orientation.

### Construction of elongated silicatein-inducible strains

In order to engineer bacterial microlenses with enhanced asymmetrical shapes, hyper-elongated *E. coli* cells were generated through both genetic engineering and small-molecule treatment approaches. Inducible overexpression of SulA from a plasmid or the addition of the drug cephalexin resulted in cells that adopted long, filamentous cell shapes (**Supplemental Figure 11, Supplemental Table 1**). The effect of SulA overexpression on cell length over time following induction was measured by fixing the cells at different timepoints and assessing cell morphology via light microscopy. Cells expressing SulA were significantly longer than unmodified cells or uninduced cells at every timepoint in the induction process, with SulA cells at the 6-hour timepoint reaching the longest observed length of 15.3 α 4.4 µm, in contrast to 1.6 α 0.3 µm for wild-type cells (**Supplemental Figure 12A, B**).

To create elongated cells that can encapsulate themselves in polysilicate, SulA overexpression was induced simultaneously with either TaSil or SdSil overexpression (TaSil+SulA or SdSil+SulA) for 3 hours, after which SulA inducers were removed to limit elongation, and cultures were incubated with orthosilicate for 3 additional hours. Control cultures were induced for SulA overexpression, but silicatein overexpression was not induced. Quantification of the lengths of the cells indicated that TaSil+SulA cells without silicatein induction grew longer than SulA cells but displayed less elongation upon silicatein induction, with co-induced TaSil+SulA cells reaching 13.7 α 3.9 µm by the end of the 6-hour encapsulation procedure in comparison with 17.9 α 5.6 µm for the TaSil+SulA cells without silicatein induction (**Supplemental Figure 12A, C**). SdSil+SulA cells without silicatein induction became less elongated than the uninduced SulA cells, reaching 9.9 α 2.4 µm. Silicatein induction caused the SdSil+SulA cells to become somewhat more elongated, though on average shorter than the TaSil+SulA cells, showing lengths of 13.6 α 3.6 µm at 6 hours (**Supplemental Figure 12A, D**).

Cephalexin treatment of wild-type cells (WT+Ceph) resulted in cell lengths of 6 α 1.7 µm at 6 hours, which represents significant elongation in comparison to wild-type cells but less elongation than for any of the SulA-overexpressing strains (**Supplemental Figure 13A, B**). To evaluate conditions for polysilicate-encapsulation of these drug-induced elongated cells, silicatein inducers and cephalexin were added to cultures of TaSil or SdSil (TaSil+Ceph or SdSil+Ceph) at the same time, and after three hours of incubation cephalexin was removed and the cells were incubated with orthosilicate for an additional 3 hours. TaSil+Ceph and SdSil+Ceph cells were significantly more elongated than unmodified or uninduced cells, with induced TaSil+Ceph becoming more elongated (7.8 α 2.3 µm) (**Supplemental Figure 13A, C**) than induced SdSil+Ceph cells (4.1 α 1 µm) (**Supplemental Figure 13A, D**). Overall, we observed that the genetically-induced cell elongation treatments resulted in more highly elongated cells than the small-molecule treatments for all conditions.

### Encapsulation of elongated cells in polysilicate

To analyze polysilicate localization patterns of elongated silicatein-expressing cells, strains were induced for silicatein expression simultaneously with induction of SulA expression or addition of cephalexin, followed by removal of inducer molecules and incubation with orthosilicate. Cells were stained with Rhodamine123, and polysilicate localization was visualized via confocal fluorescent microscopy. For SulA-expressing strains, SdSil+SulA cells showed Rhodamine123 staining that was localized to the outer borders of the elongated cells (**Figure 4A**), in comparison to TaSil+SulA cells that showed staining of internal inclusion bodies and the non-silicatein-expressing SulA cells that displayed a more diffuse, dimmer staining pattern (**Figure 4B and C**). The SdSil+SulA and TaSil+SulA cells both had a ratio of border-to-cytoplasm fluorescence signal that was significantly higher than for the non-silicatein-expressing SulA cells, which had a ratio close to 1 (**Figure 4D**). For the cephalexin-treated cells, the TaSil+Ceph and SdSil+Ceph cells both showed border-localized Rhodamine123 staining patterns (**Figure 4E and F**), while the cephalexin-treated wild-type cells showed a more diffuse staining pattern with some slight cell membrane staining (**Figure 4G**). The TaSil+Ceph and SdSil+Ceph cells showed a significantly higher ratio of border-to-internal fluorescence staining than the WT+Ceph cells (**Figure 4D**).

**Figure 4:**
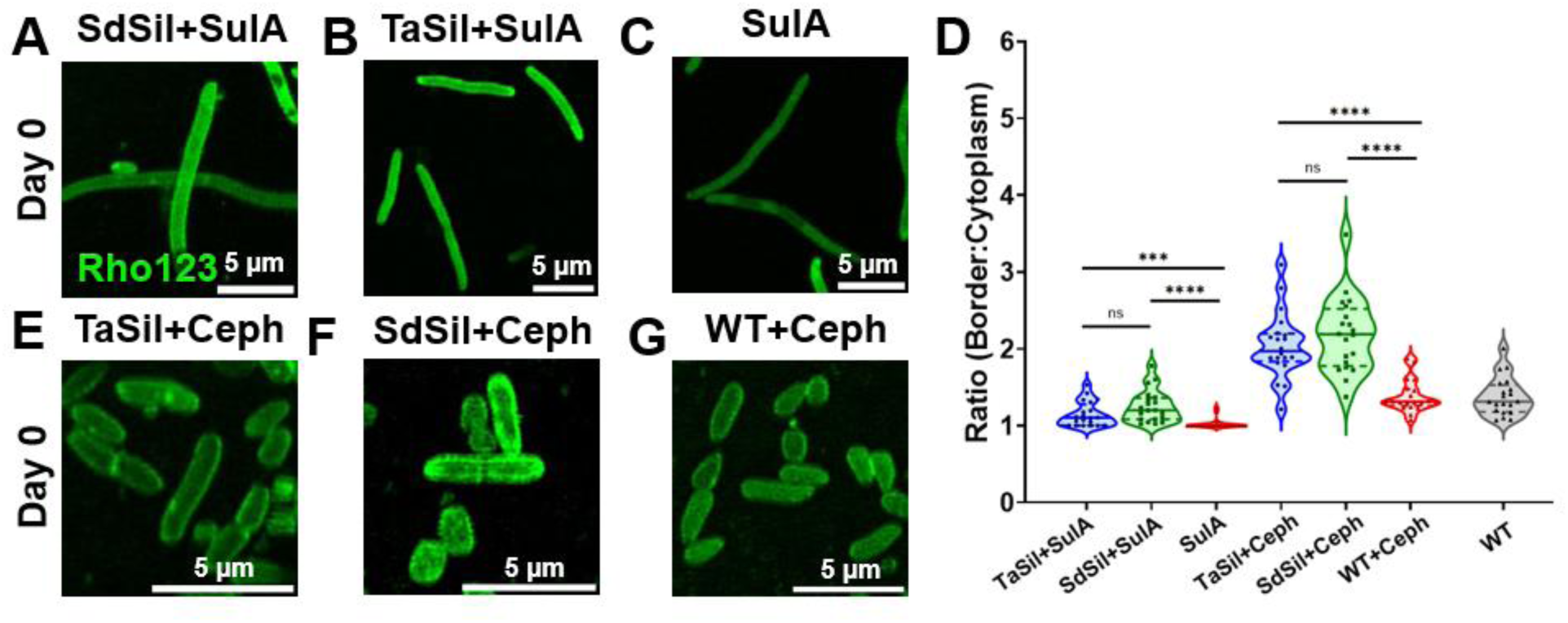
Polysilicate localization of highly elongated, polysilicate-encapsulated bacterial cells. (A-C and E-G) Cells stained with Rhodamine123 at day 0 where (A) are SdSil+SulA, (B) is TaSil+SulA, (C) is SulA, (E) is TaSil+Ceph, (F) is SdSil+Ceph, and (G) is WT+Ceph. (D) The quantification of the ratio of Rhodamine123 border to cytoplasm staining for strains A-C and E-F and WT. (n=20) ns: no significance,*** p≤0.001, **** p≤0.0001.

To determine the pattern of polysilicate localization on elongated cells over time, Rhodamine123 staining and confocal fluorescence microscopy were performed at regular monthly intervals on cells that were stored in buffer. Stored SdSil+SulA cells maintained elongated cell shapes with Rhodamine123 staining at their outer borders that was enhanced relative to freshly prepared samples (**Figure 4A**) at timepoints from 1 month (**Supplemental Figure 14A**) through 6 months (**Supplemental Figure 14B**). Both SdSil+SulA and SulA control cells showed the appearance of dispersed internal inclusion bodies at the later timepoints (**Supplemental Figure 14A-D**), while TaSil+SulA cells showed inclusion bodies at all timepoints that increased in abundance over time (**Supplemental Figure 14E-G**). The cephalexin-treated cells displayed less intense border staining over time for both TaSil+Ceph (**Supplemental Figure 14G-I**) and SdSil+Ceph cells (**Supplemental Figure 14G, J, and K**), while the WT+Ceph control cells showed slight increases to border staining at 1 month (**Supplemental Figure 14G and L**) followed by decreased border staining at subsequent timepoints (**Supplemental Figure 14G and M**).

To monitor the maintenance of the altered shapes over time for the elongated, polysilicate-encapsulated cells, the lengths of the cells were measured at regular monthly intervals. The SdSil+SulA cells and SulA control cells were observed to significantly decrease in length over time (**Supplemental Figure 14N**), likely due to the lack of exposure to the SulA inducer molecule during storage allowing the Z-rings to complete cell division(s). The TaSil+SulA cells were not as elongated as the other SulA-expressing cells and displayed no significant changes in length at subsequent timepoints (**Supplemental Figure 14N**). The cephalexin-treated cells all showed minimally increased cell length throughout the experiment relative to unaltered wild-type cells, with cell lengths around 1-3 μm for all timepoints and strains (**Supplemental Figure 14N**). Since the cephalexin-treated cells were less elongated at all timepoints than the SulA-expressing cells, and the TaSil+SulA strain exhibited inclusion bodies as well as less elongation upon silicatein induction, these strains were not utilized further in this work.

To analyze the effect of polysilicate encapsulation on the morphology of elongated cells, thin sections were prepared and imaged via TEM of SdSil+SulA cells that were co-induced for silicatein and SulA expression and then incubated with orthosilicate, as well as untreated wild-type cells and SulA cells that were induced for SulA expression. The polysilicate-encapsulated SdSil+SulA cells showed cell borders with smooth morphologies (**Supplemental Figure 15A**), similar to the smooth cell surfaces seen previously for polysilicate-encapsulated cells^1^. The non-encapsulated SulA and wild-type cells displayed the ruffled surfaces typically seen for *E. coli* cells^18^ (**Supplemental Figure 15B and C**). Quantifications of the TEM images indicated that the polysilicate-encapsulated SdSil+SulA cells had significantly lower surface roughness as well as significantly less electron-dense interiors compared to either the SulA or the wild-type non-encapsulated control cells (**Supplemental Figure 15D and E**). SEM imaging of the elongated strains indicated that the polysilicate-encapsulated SdSil+SulA cells appeared more highly embedded under the high-density imaging conditions than the non-encapsulated SulA or wild-type cells (**Supp. Figure 16**). These experiments demonstrated that the SdSil+SulA cells were able to encapsulate themselves in a layer of polysilicate while also adopting a highly elongated cellular morphology, both of which were maintained for at least six months.

### Elongated, polysilicate-encapsulated cells scatter focused light across long distances

Since rod-shaped *E. coli* cells that are encapsulated in polysilicate show light-focusing properties that are dependent on their orientation, we hypothesized that the elongated, polysilicate-encapsulated cells would display heightened orientation-dependence of their light-scattering. Both SdSil+SulA and SulA cells are highly elongated, with aspect ratios of 8 α 2.8 for polysilicate-encapsulated SdSil+SulA cells and 6.5 α 1.8 for SulA cells, compared to 2 α 0.3 for polysilicate-encapsulated SdSil cells and 1.5 α 0.3 for wild-type *E. coli* cells, which are both rod-shaped (**Supplemental Figure 17**).

MAIM microscopy was applied to analyze the light-scattering properties of elongated, polysilicate-encapsulated SdSil+SulA cells in comparison to rod-shaped polysilicate-encapsulated SdSil cells, elongated non-encapsulated SulA cells, and rod-shaped non-encapsulated wild-type cells. Maximum intensity projections showed that SdSil+SulA cells and SulA cells both scattered bright jets of light similar to that of the SdSil cells, whereas the wild-type cells did not (**Supplemental Figure 18A**). Quantification of the intensity of the scattered light as a function of distance from the cells revealed that the elongated SdSil+SulA and SulA strains both scattered light with a similar maximum intensity that was more than twice as high compared to the rod-shaped SdSil cells (**Supplemental Figure 18B**). SdSil+SulA cells showed a focal peak located approximately 2.5 μm away from the cells, whereas the light intensity for the SulA cells was at a maximum within the first micrometer from the cells and decreased with increasing distance. The length of the scattered light was not significantly different between SdSil+SulA cells and either SdSil or SulA cells but was significantly higher than for wild-type cells (**Supplemental Figure 18C**). The width of the scattered light was not significantly different between SdSil+SulA cells and SulA cells, though it was significantly higher than for SdSil and wild-type cells (**Supplemental Figure 18D**). The integrated intensity of the scattered light was significantly higher for SdSil+SulA cells compared to any of the other strains (**Supplemental Figure 18E**), and the brightness of the cell bodies for the elongated SdSil+SulA and SulA cells were significantly higher than for either the rod-shaped SdSil and wild-type cells (**Supplemental Figure 18F**). These data indicate that the elongated cells were all able to scatter jets of light, with the polysilicate encapsulation of the elongated SdSil+SulA cells resulting in an enhanced total amount of scattered light and the appearance of a focal peak that was twice as bright as the rod-shaped, polysilicate-encapsulated cells.

To analyze the effect of cell orientation on light scattering of the elongated cells, the SdSil+SulA and SulA cells were divided into bins based on the angle of the long axis of the cell relative to the incident light. Maximum projections of individual cells showed that both elongated, polysilicate-encapsulated SdSil+SulA cells and elongated, non-encapsulated SulA cells in the 0-30° bin scattered intense jets of light, while cells in the 31-60° bin scattered weaker jets of light emanating from the cell poles, and cells in the 61-90° bin scattered low-intensity light from the cell poles (**Figure 5A and B**). Scattering profiles indicated that the SdSil+SulA cells in the 0-30° bin scattered light with the highest intensity across the entire scattering range, with a sharp focal peak 1.95 µm from the cell, while cells in the 31-60° and 61-90° bins scattered light with increasingly lower intensity and broader focal peaks (**Figure 5C and D**). The SulA cells also showed decreasing intensity of scattered light for cells that were positioned at higher angles, with none of the bins showing prominent focal peaks (**Figure 5C and E**). Apart from the dramatic effects on scattering profiles, the angle of the cells was determined to have little measured effect on the length or width of the scattered light, or for the integrated light intensity for the scattered light or for the cell bodies (**Supplemental Figure 19**).

**Figure 5:**
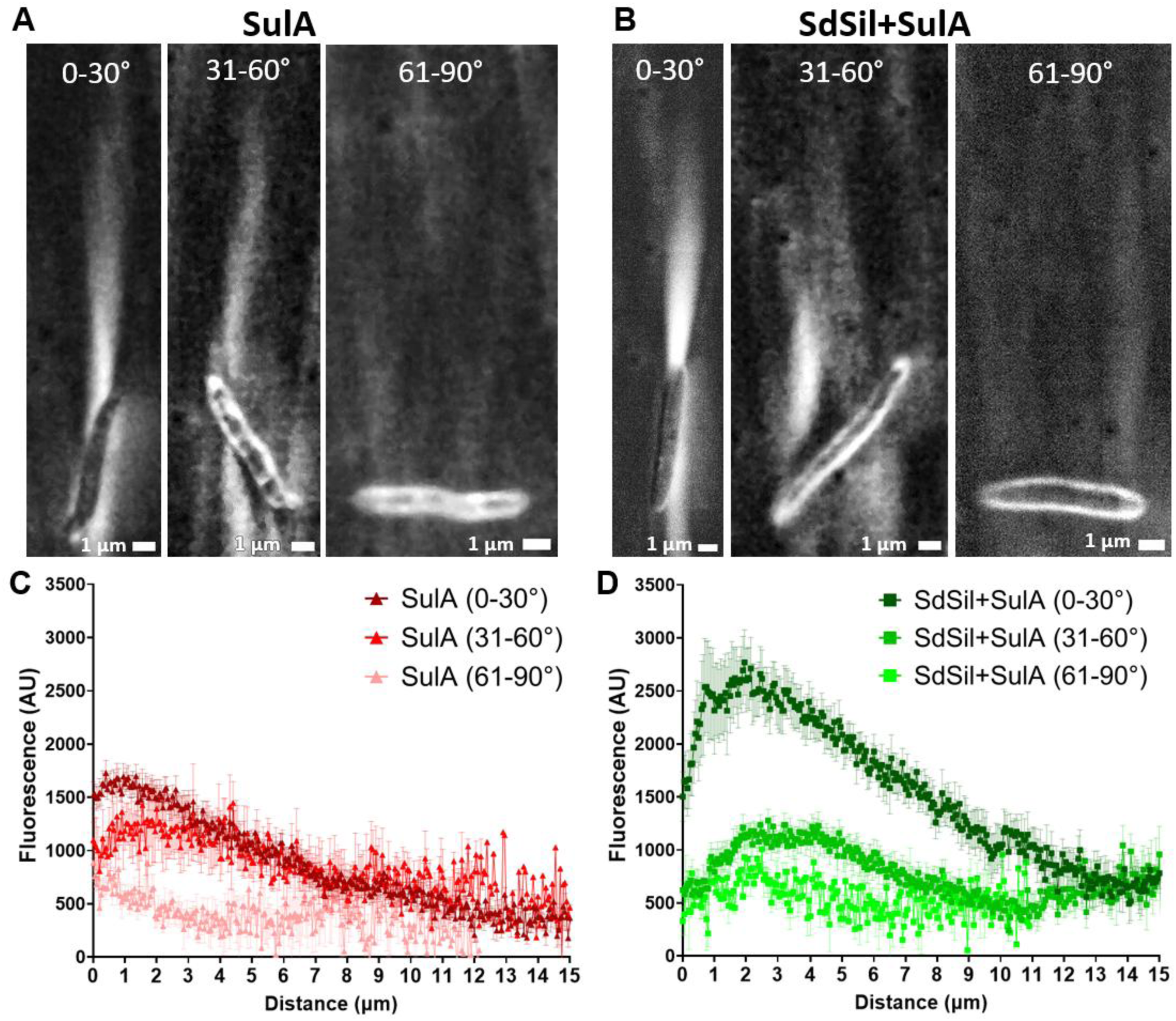
Light scattering abilities of elongated, polysilicate-encapsulated cells. (A and B) Maximum intensity projections for (A) elongated, non-encapsulated SulA and (B) elongated, polysilicate-encapsulated SdSil+SulA cells scattering light via MAIM for cells binned into 0-30°, 31-60°, and 61-90° relative to the incident light. (C-D) Profiles of the scattered light as a function of the distance from the edge of the cell, calculated from the maximum intensity projections, where error bars correspond to standard error of the mean, for (C) SulA and (D) SdSil+SulA. (n_SulA,0-30°_=19, n_SulA,31-60°_=4, n_SulA,61-90°_=6, n_SdSil+SulA,0-30°_=9, n_SdSil+SulA,31-60°_=18, n_SdSil+SulA,61-90°_=3)

Overall, these data show that the light scattered by elongated, polysilicate-encapsulated cells shows heightened orientation dependance relative to cells with unmodified rod shapes. The elongated cells primarily scatter light from their rounded poles, with little light scattered from the mid-sections of their cell bodies. Elongated cells that are oriented nearly parallel to the incident light are able to scatter the light with the highest intensity observed for any of our engineered cells. The focal peak of this scattered light is located 2-3 μm away from the distal edge of the bacteria cells, which can be tens of micrometers away from the proximal edge of the elongated cells, representing transmission of the input light through the cell bodies along distances of 10 μm or farther.

### Polysilicate-encapsulated, shape-altered cells maintain metabolic activity

To determine how the physiology of the engineered cells was affected by polysilicate encapsulation in combination with shape alterations, their growth and metabolic activity were evaluated. Growth curves were measured for the cells during induction of silicatein and shape alteration followed by polysilicate encapsulation. For the BolA- and SulA-expressing cells, the co-induction of shape-alteration and silicatein resulted in stalling of cell growth beginning around the time of orthosilicate addition (**Supplemental Figure 20A and B**). A22-treated wild-type cells exhibited accelerated growth compared to untreated wild-type cells, with A22-treated silicatein expressing cells exhibiting similar growth kinetics to cells expressing only silicatein (**Supplemental Figure 20C**). The effect of polysilicate encapsulation combined with shape alteration on bacterial cell division capabilities over time was measured by a colony forming unit (CFU) assay. Initially, spherical polysilicate-encapsulated strains SdSil+BolA and TaSil+A22, as well as the elongated polysilicate-encapsulated strain SdSil+SulA, all showed a significant reduction in CFU/mL values compared to wild-type cells, with the spherical SdSil+BolA and TaSil+A22 both showing approximately 2-log reductions while the elongated SdSil+SulA showing greater than 5-log reduction (**Figure 6A, B**). Following storage, CFU/mL values decreased over time for all strains, reaching undetectably low levels after 5 months for SdSil+BolA and after 4 months for SdSil+SulA, each of which was one month earlier than for their respective non-silicatein-expressing control strains BolA and SulA (**Figure 6A, B**). The TaSil+A22 cells showed an approximate 5-log decrease in CFU/mL during 6 months of storage, but these cells still retained substantial cell-division capability after 6 months, reflecting the extended cell-division lifespan observed for the A22-treated wild-type cells (**Figure 6A**). These results indicate that the shape-altered, silicatein-expressing cells have greatly reduced capacity for cell division following encapsulation.

**Figure 6:**
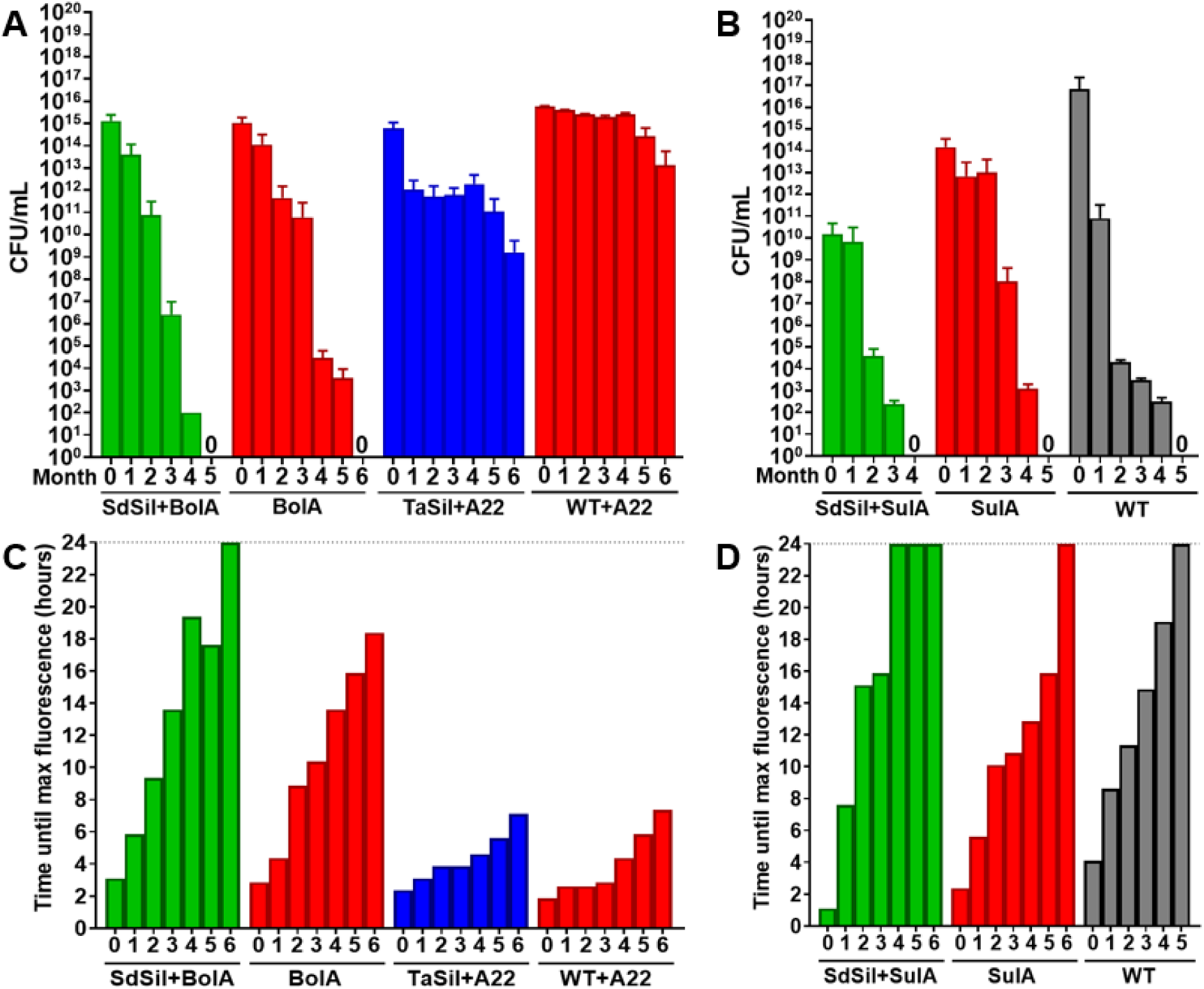
Shape-altered, encapsulated cells have reduced cell division but retain metabolic activity. (A and B) Colony forming unit assays for (A) spherical cells and (B) elongated cells following induction of shape-alteration and silicatein expression and polysilicate-encapsulation over 6 months of storage. (C and D) Time until maximum fluorescence for alamarBlue metabolic assays for (C) spherical cells and (D) elongated cells following induction of shape-alteration and silicatein expression and polysilicate-encapsulation over 6 months of storage. The dashed line at 24 hours for C and D corresponds to the end of the 24-hour experiment, where data collection ended.

The cells’ metabolic activities were measured over time using the cell viability reagent alamarBlue to detect the ability of the cells to take up and metabolically reduce resazurin molecules to the fluorescent compound resorufin^19^. Previously, post-encapsulated rod-shaped cells showed no detectable metabolic activity by 4-5 months^1^. Shape-altered, spherical encapsulated cells maintained metabolic activity for 5 months or longer, while the elongated encapsulated cells showed metabolic activity for 3 months (**Figure 6C and D**). The time to reach peak fluorescence increased with incubation time for all strains, and the A22-treated cells maintained levels of metabolic activity that were higher than the other strains over the time course of the experiment (**Figure 6C and D**). Overall, these experiments indicate that the shape-altered, polysilicate-encapsulated cells demonstrate metabolic activity for several months post-encapsulation, despite showing lowered cell division capability.

### Heat killing encapsulated cells does not compromise their light-scattering abilities

Since our rod-shaped and spherical polysilicate-encapsulated bacterial cells scattered robust light and maintained considerable metabolic and divisional activity for several months post-encapsulation^1^, we investigated how this scattering would be affected by purposeful heat inactivation of the cells. Heat inactivation was most successful when the strains were exposed to 75°C for one hour. Following this treatment, all strains were observed to be no longer metabolically active, as determined using the alamarBlue assay, and no longer divisionally active, as determined using the CFU assay (**Supplemental Figure 21A and B**). The heat-killed cells showed similar Rhodamine123 staining patterns to their live counterparts (**Supplemental Figure 21C**). The heat-killed polysilicate-encapsulated strains (TaSil, SdSil, SdSil+BolA, and TaSil+A22) all displayed brighter signals at the cell borders than internally, though with a more punctate pattern than the live cells (**Supplemental Figure 21C and D**). The heat-killed control strains (WT, BolA, and WT+A22) maintained the diffuse, lower-intensity staining pattern seen for the live cells, though intensity did increase due to cell death^20^ (**Supplemental Figure 21C and D**).

To determine whether purposefully inactivated cells maintain their light-scattering properties, we performed MAIM experiments on the heat-killed cells. The heat-killed cells were observed to scatter light (**Supplemental Figure 22**) that was as bright or brighter than the corresponding live cells (**Supplemental Figure 23A-D**). For the rod-shaped cells, heat-killing caused the polysilicate-encapsulated cells to scatter light of equivalent intensity with a focal peak approximately 1.8 µm from the cells for TaSil (TaSil HK), or higher intensity light with a focal peak approximately 2.2 µm from the cells for SdSil (SdSil HK) (**Supplemental Figure 23A**). The jets of light scattered by rod-shaped, heat-killed cells were slightly shorter and narrower than for their live counterparts (**Supplemental Figure 23E and F**).

For the spherical, BolA-expressing cells, the heat-killed cells scattered light with similar intensity as the live SdSil+BolA strain, where the polysilicate-encapsulated cells (SdSil+BolA HK) had a focal peak approximately 2.7 µm from the cells (**Supplemental Figure 23B**). The heat-killed BolA-expressing cells scattered jets of light with equivalent length and width to the live cells (**Supplemental Figure 23E and F**). The heat-killed, spherical, A22-treated cells also scattered focused light, which was higher intensity than the live A22-treated cells (**Supplemental Figure 23C**). A22-treated, encapsulated cells did not scatter significantly different amounts of light than the non-encapsulated cells, even when heat killed. The scattered light from the heat killed, A22-treated showed similar length and width to the live, A22-treated cells (**Supplemental Figure 23E and F**).

These data indicate that the purposeful sterilization of bacterial microlenses through heat treatments does not diminish their light scattering and focusing abilities. This treatment would enable the use of these bacterial microlenses in applications without fear of microbial escape, since they retain full optical functionality despite the inactivation of their cellular physiology.

## Discussion

In this work, we have described the creation of both spherical and elongated bacteria encapsulated in polysilicate, and we have demonstrated that these different morphologies alter the interaction with light in a controlled manner. Through overexpressing the BolA gene or treating with the drug A22 while displaying the sea-sponge silicatein enzyme on the surface of *E. coli* cells, we were able to create spherical polysilicate encapsulated bacterial cells that act as microlenses and scatter more intense, focused light compared to the previously reported rod-shaped polysilicate-encapsulated bacterial cells, without orientation dependence. In contrast, through overexpressing the SulA gene while displaying the silicatein enzyme, we have created highly elongated polysilicate-encapsulated bacterial cells that also scatter more intense, focused light compared to the rod-shaped polysilicate-encapsulated bacterial cells, in a highly orientation-dependent manner. Their elongated shape has light-guiding properties, where encapsulated cells within the 0-30° tilt range display a significant increase in scattered light emitted at or very near the exiting tip of the cell. This behavior is akin to light trapped within a photonic waveguide or fiber optic cable. Both of these micron-sized optical devices are metabolically active for months following the shape-alteration and polysilicate-encapsulation process. These photonic devices with engineered optical properties were created solely through the manipulation of the bacterial cell itself, without the need for machining or mechanical manipulations. We have therefore demonstrated the creation of self-assembled bacterial photonic devices with a broad range of tunable optical properties, spanning applications from microlenses to waveguides.

Additional tuning of the size and shape of *E. coli* is possible using a variety of techniques. For instance, the mean length of *E. coli* cells can be finely calibrated by changes in the pH of the growth medium^21^. Our results also emphasize that bio-engineered photonic devices derived from cells are not limited to the shapes found in wild-type cells. Using insights gained by previous research into cell shape regulation, even more structures should be achievable. In addition to genetic and pharmacological manipulation of cell shape, it has been shown that *E. coli* can be mechanically manipulated to adopt different morphologies. *E. coli* can invade narrow channels by altering its cell morphology into a flattened disc^22^. Using PDMS molds, filamentous bacteria can be formed into spiral, crescent, and sinusoid shapes^14^. By combining PDMS molds with drugs that inhibit cell wall formation, *E. coli* can be sculpted into square, triangular, and other arbitrary geometries^11^, each of which are expected to exert different effects on the cells’ optical properties.

While many potential techniques exist to manipulate the shape of *E. coli* cells, an even wider design space of bio-engineered photonic devices can be explored by using other species as a chassis organism for biological engineering. Various species of bacteria can exhibit a broad range of sizes and shapes not easily achieved in *E. coli*. The smallest reported bacteria can have diameters less than 0.2 µm^23^, while the largest cocci bacteria can have diameters of over 100 µm^24^ and the longest rod-shaped bacteria have been reported to grow to over a centimeter in length^25^. However, many of these species are difficult to culture or difficult to genetically manipulate; it will be important to identify strains with unique size and shape characteristics that are also able to be bio-engineered to display biomineralization enzymes. Moving beyond bacteria, other model organisms can also provide interesting size and shape possibilities. Yeast cells are typically ~10x longer than *E. coli* and also exhibit a range of morphologies that include both round and filamentous cells^26^. Additionally, neurons from zebrafish have been modified so that the filamentous axons can be steered to grow in precise locations using optogenetics, providing a possible way to engineer waveguide connections between specific points^27^.

We anticipate a range of applications for both round and filamentous living, nanophotonic particles produced in a sustainable and low-cost manner using bacteria. We show that round bacteria are more effective at focusing light and do not require alignment, making them easier to incorporate into larger devices than rod-shaped cells. The small size of spherical bacterial microlenses makes them well suited for improving the efficiency of miniaturized photodetectors like those used in mobile phones, which now require pixel sizes approaching the submicron scale^28^. Bacteria have been shown to spontaneously pattern themselves on periodic nanostructures^29^, making it possible to autonomously position bacterial microlenses over the pixels of a photodetector. Filamentous bacteria are less well suited as microlenses but are promising candidates for forming nanophotonic waveguides. Nanophotonic waveguides made of living cells have been used as sensitive and biocompatible biosensors^30^, and a chain of individual bacteria trapped between two fiber ends has previously been shown to act as a waveguide and biosensor^31^. A similar approach could be used to immobilize the elongated waveguide cells in a fixed orientation, allowing for integration of these cells into a biosensing device. A single elongated cell would be expected to provide a sturdier structure for guiding light than a chain of individual cells held together by optical forces. A single-cell waveguide would also avoid stochastic variations in gene expression, and potentially optical properties, due to the bursty dynamics of mRNA production in individual cells^32^, providing more uniform behavior across the device. Further study is needed to compare the light-guiding performance of single elongated cells versus chains of multiple smaller cells, as well as to determine the degree to which a polysilicate coating improves the efficiency of a single-cell waveguide-based biosensor.

## Methods

### Strain information

The strain background used for all experiments is Top10 *Escherichia coli* (genotype: F- mcrA Δ(mrr-hsdRMS-mcrBC) φ80lacZΔM15 ΔlacX74 recA1 araD139 Δ(araleu) 7697 galU galK rpsL (StrR) endA1 nupG), which is a non-motile laboratory strain. TaSil and SdSil plasmids were designed and cloned as previously described^1^. BolA (DNA-binding transcriptional dual regulator BolA, EcoCyc EG10125) was placed behind a strong ribosome binding site (B0034^33^) and amplified with adaptamers containing restriction sites KpnI and PstI from iGEM part BBa_K1890030^34^, then cloned into the multiple cloning site of pBAD33^35^ using KpnI and PstI, such that control over its expression was regulated by an arabinose operon. SulA (EcoCyc EG10984) was placed behind the same ribosome binding site (B0034) and contiguous to the arabinose operon (araC, araBAD promoter) from the SulA pBAD33 plasmid. This cassette was cloned into the multiple cloning site of pSB1C3^36^using EcoRI and XbaI, placing SulA under control of the arabinose operon in a different plasmid backbone. Each silicatein plasmid (TaSil or SdSil) was co-transformed into Top10 cells along with either the BolA pBAD33 or SulA pSB1C3 plasmid so that the constructs could be co-induced. **Supplemental Table 2** summarizes the constructs, their resistances, and inducers.

### Cell orientation binning

Using the TaSil and SdSil datasets from Sidor *et al.* PNAS (2024)^1^ the MAIM images were reanalyzed to determine the angle of the cell in relation to the incident light. Cells were divided into three bins: 0-30°, where the long axis of the cell was parallel with the incident light; 31-60°, where the cells were diagonal to the incident light; and 61-90°, where the cells were perpendicular to the incident light (see **Figure 1A**). The TaSil and SdSil datasets were pooled together, and the profiles of the scattered light were determined for the three cell orientation bins. Using GraphPad, the area under the curve function was used to determine the focal peak values for each angle bin. The same data sets were analyzed to determine the length and width of scattered light and the integrated density of the scattered light graphs.

### Culture growth conditions

Cell growth was performed as previously described^1^ with the additions of induction steps for BolA and SulA or addition of drugs S-(3,4-dichlorobenzyl)isothiourea (A22) (Sigma Aldrich, CAS: 475951) or the β-lactam antibiotic cephalexin (CAS: 1820673-23-1). Briefly, overnight cultures were prepared for each strain, and the next morning fresh subcultures were prepared. Cells were grown in Luria-Bertani (LB) media (plus antibiotics – see **Supplemental Table 2**; ampicillin for silicatein plasmids, and chloramphenicol if using BolA or SulA plasmids), in liquid suspension in an Erlenmeyer flask, at 37°C for 2 hours with constant shaking at 250 rpm. After 2 hours, silicatein was induced (see **Supplemental Table 2**; IPTG for TaSil and rhamnose for SdSil), and shape change was induced. For BolA and SulA induction, a final concentration of 30 mM arabinose was added to the cultures at the same time as the TaSil and SdSil induction.

For A22, a final concentration of 10 µg/mL A22 was added to the cultures at the same time as the TaSil and SdSil induction. For cephalexin, 25 µg/mL of cephalexin was added to the cultures at the same time as the TaSil and SdSil induction. Cells were grown for 3 hours at 37°C with constant shaking at 250 rpm. For SulA and cephalexin cells, after 3 hours of incubation with the inducers/drug, cells were pelleted at 4500 rpm for 10 minutes in a swinging bucket centrifuge (Thermo Scientific, Sorvall ST 16R). The supernatant containing the inducers/drugs was removed, and cells were resuspended in fresh LB and antibiotics, as well as fresh silicatein inducers, in order to remove the arabinose or cephalexin exposure so that cells grew to desired lengths. Sodium orthosilicate (Na_4_SiO_4_, Alfa Aesar, CAS: 13472-30-5) was then added to a final concentration of 100 µM sodium orthosilicate for each silicatein-expressing culture. Cells were incubated with the sodium orthosilicate at 37°C with constant shaking at 250 rpm for 3 hours.

After 3 hours, cells were pelleted at 4500 rpm for 10 minutes in a swinging bucket centrifuge. Cells were washed 3 times with 1X Tris buffered saline (TBS: 50 mM Tris-HCl, 150 mM NaCl, pH 7.5) and resuspended in 1X TBS and stored at 4°C with constant rotation (Benchmark Scientific Roto-Mini Plus Variable Speed Rotator R2024) at 30 rpm until use.

### Cell fixation for encapsulation protocol evaluation

During the encapsulation protocol described above, at one-hour timepoints beginning at the addition of the inducer molecules and/or drugs, an aliquot of 1 mL of cells was fixed in 2.5% glutaraldehyde in 1X phosphate buffered saline (PBS – MP Biomedicals, CAS: 2810306) for 1.5 hours at room temperature. Cells were washed three times with 1X PBS and resuspended in 1X PBS. Fixed cells were stored in 1X PBS at 4°C with constant rotation (Benchmark Scientific Roto-Mini Plus Variable Speed Rotator R2024) at 30 rpm until imaged using MAIM microscopy. Transmitted light illumination was used to image the fixed cells, without the addition of stains or dyes.

### Circularity calculations

The circularity of bacteria cells was calculated using FIJI software^37^. For each sample image, the image was thresholded to segment the cell boundaries. For the cells in each thresholded image, the Analyze Particles tool was applied to measure and report the perimeter, area, length, and width of the cell. Circularity was then calculated using: 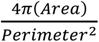.

### Cell length and width analyses

Using FIJI, a line was drawn lengthwise along the cell to determine the longest axis of the cell, which was defined as the length of the cell. A second line was drawn widthwise to determine the shortest axis of the cell, which was defined as the width of the cell.

### Rhodamine123 staining

Rhodamine staining was performed as previously described^1^. Briefly, after bacteria culture growth, shape change, and incubation with sodium orthosilicate, live cells were stained with Rhodamine123 (Invitrogen R302). 1/10 volume of 500 µm Rhodamine123 was added, and cells were incubated for 15 minutes in the dark at room temperature. Cells were washed three times with 1X TBS and resuspended in 1X TBS before being pipetted onto a 1% agarose pad and sealed with a coverslip. Cells were imaged on a Nikon A1R HD Laser Scanning Confocal Microscope using a 60X oil Apochromat TIRF objective (1.49 NA). Rhodamine123 signal was imaged using a 488 nm excitation laser and a 500-550 nm emission filter. Images were analyzed using FIJI software.

### Cell preparation for Transmission Electron Microscopy (TEM)

Following growth and induction with chemical inducers and sodium orthosilicate, cells were fixed overnight at 4 °C with continuous rotation in a solution containing 2.5% glutaraldehyde prepared in 0.1 M sodium cacodylate buffer. After fixation, the cells were pelleted and washed twice for 10 minutes each using the same buffer. Post-fixation, samples were treated with 1% osmium tetroxide for 30 minutes. The cells were again centrifuged and rinsed twice with deionized water for 10 minutes per wash, after which the supernatant was discarded.

The resulting cell pellets were embedded in 3% agarose, and the solidified agarose was cut into approximately 1 mm cubes. These cubes were dehydrated through a graded ethanol series (50%, 65%, 80%, 95%, and 100%), with each step lasting 20 minutes, and the 100% ethanol step was repeated three times. The dehydrated samples were then placed into a 1:1 mixture of propylene oxide and ethanol for 30 minutes, followed by two additional 30-minute incubations in pure propylene oxide. Samples were incubated with a 1:1 mixture of epoxy resin and propylene oxide under rotation for 2.5 hours, then transferred to 100% epoxy resin and left overnight.

Polymerization was carried out at 65 °C for 48 hours in embedding molds. Ultrathin sectioning was performed using a Leica UC7 ultramicrotome with a diamond knife, first generating 1 µm sections. These samples were stained on glass slides with toluidine blue, rinsed with deionized water, and inspected by light microscopy to identify regions for thin sectioning. Selected areas were sectioned to 70 nm thickness and mounted on formvar/carbon-coated nickel slot grids (Electron Microscopy Sciences, Hatfield, PA). Grids were subsequently stained with 2% aqueous uranyl acetate followed by 3% lead citrate. Imaging was conducted using a Hitachi 7650 transmission electron microscope operating at 80 kV, equipped with a Gatan Erlangshen 11-megapixel digital camera.

### Cell surface roughness calculations

Surface roughness of cells was determined as previously described^1^. Using FIJI, individual cells were selected, and the cell images were duplicated. A 2-pixel Gaussian blur was performed on one duplicate, and a 10-pixel Gaussian blur was performed on the other. Using the lasso tool, the outer edge of the cell was outlined, and the perimeter was measured for both blurred images. The cell surface roughness was calculated as: Perimeter_(Blur2)_ – Perimeter_(Blur10)_.

### Scanning electron microscopy (SEM)

Samples stored in 1X TBS buffer were centrifuged, pelleted, and washed twice with distilled water. For SEM analysis, approximately 100 µL of each sample was deposited onto an SEM sample holder and dried overnight. The dried samples were sputter coated with a 5 nm layer of platinum to enhance conductivity. SEM imaging was performed using a Hitachi SU3500 in secondary electron mode with an accelerating voltage of 10 keV and a working distance of 5 mm.

### Microscope set-up

MAIM was performed using a microscope set-up as described in Sidor, *et. al*. PNAS (2024)^1^. A custom-built inverted fluorescence microscope was used featuring a high NA objective (Nikon CFI Apochromat TIRF 100XC Oil). Fluorescence illumination was delivered by a 488 nm laser beam that was coupled through a single mode fiber and then directed through the objective via a dichroic mirror. A motorized translation stage attached to the fiber launcher allowed the user to translate the position of this beam and thereby vary the angle of illumination on the sample from 0° (epi illumination) down to +/- 90° (total internal reflection). Fluorescent excitation collected through the objective then passed through the dichroic mirror and was imaged onto a sCMOS camera (Thorlabs).

### Multiple angle illumination microscopy (MAIM)

MAIM imaging was performed as described in Sidor, *et. al*. PNAS (2024)^1^. Briefly, an agarose pad was prepared by melting 1% agarose in a 0.01% poly-lysine solution and solidifying it into a 7/16 diameter circle (0.5 mm thick, ~100 µL in volume). The agarose pad was stained for 20 minutes with 20 µL of 0.5 mg/mL Alexa Fluor™ 488 NHS Ester (Invitrogen A20000) dissolved in a solution of 0.1 M sodium bicarbonate pH 8.3 in a dark humidity chamber. Excess dye was quenched with 1 M glycine rinsed over the agarose pad. 10 µL of cells was pipetted onto the stained agarose pad and sealed with a cover slip. Cells were imaged by a custom-built inverted fluorescence microscope (see Microscope set-up). The angle of illumination was continuously varied from −90° to +90°, spanning all possible illumination angles, while recording the fluorescence on the camera at 30 fps. The laser intensity was chosen to be relatively low to minimize photodamage (200 µW total, ~0.005 µW/µm^2 in the image plane).

### MAIM image processing

MAIM image processing using FIJI was performed as previously described^1^ with slight changes for the following parameters: integrated density and cell body mean gray value. Briefly, A 50-pixel radius rolling ball background subtraction was performed on all movies, and a maximum intensity Z projection was created from each movie. The maximum intensity images were used for the following analyses: length, width, integrated fluorescence, mean fluorescence within the cell boundary, and profiles of the scattered light. The dimensions of the jet of light were defined by identifying the contiguous region downstream of the bacterium where the fluorescent intensity was above background. The length was defined by the distance from the cell edge to the far end of the jet region, the width was defined by the broadest portion of the jet region when measured perpendicular to the direction of light, and the integrated fluorescence was calculated within a rectangle the spanned the length and width of the jet region. For elongated cells, since the various angles of the cells made use of the box tool non-ideal due to extensive inclusion of non-jet regions, the lasso tool was used to determine integrated fluorescence by drawing the smallest box that included the entire jet and measuring the integrated density within that shape.

The mean light fluorescence within a cell was estimated by bounding the bacterial cell within a minimum-sized rectangle and calculating the mean intensity of the rectangle. For the elongated cells, the lasso tool was used to measure the mean gray value of the cell body, to draw the smallest box that included the entire cell and measuring the mean gray value within that shape. The scattered light profile was measured along a line through the center of the jet region starting at the cell edge and continuing to the farthest point above background.

### Aspect ratios of the cells

The following equation was used to determine the aspect ratio of the cells:

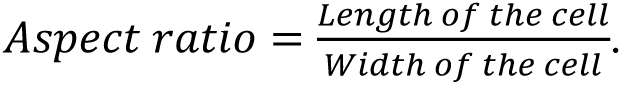

### Growth curves

Cells were cultured as described above, with the exception that cultures were grown in a 96-well plate. The plate was incubated at 37°C with continuous orbital shaking at 282 rpm in a microplate reader (BioTek Synergy H1). O.D._600_ measurements were recorded every 15 minutes for a total of 8.5 hours, with brief breaks for the addition of inducer chemical(s) and/or drugs and sodium orthosilicate. Cells were not pelleted for SulA and cephalexin growth curves, due to technical limitations inherent to using the 96-well plate set-up.

### Colony forming unit (CFU) assay

Cells were resuspended to an O.D._600_ of 0.1 in LB (with antibiotics) and were serially diluted in 1X TBS. 10 µL of the diluted cells was pipetted into a 10 µL spot onto appropriate media for each condition and dilution. Plates were incubated at 37°C overnight, and colonies were counted the following morning. CFU/mL of the samples was calculated as followed:

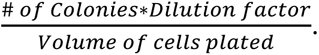

### alamarBlue cell viability assay

After growth, shape change, and incubation with sodium orthosilicate, cells were resuspended to an O.D._600_ of 0.1 in LB (with antibiotics). Cells were diluted to 10^-1^ in 1X TBS. The subsequent steps were all performed in triplicate. 100 µL of each sample was pipetted into two plates: one black, clear-bottom 96-well plate in which alamarBlue (Invitrogen DAL1025) fluorescence was recorded (Ex: 530 nm/Em: 590 nm), and one clear 96-well plate in which O.D._600_ was recorded for growth curves. Measurements were taken in separate plates because alamarBlue signal overlaps with the O.D._600_ measurements. 10 µL of alamarBlue was added to the set of samples in the black plates. Readings were taken every 15 minutes for 24 hours in a BioTek Synergy H1 microplate reader at 37°C with continuous orbital shaking (282 rpm). 1X TBS was used as a negative control for the alamarBlue assay, and LB was used for negative controls for the growth curves. The negative control values were subtracted from each experimental data set to correct for any background signals.

### Heat killing

Cells were prepared as described in the “Culture Growth Conditions” section. An aliquot of cells was taken from freshly prepared cells and diluted to an O.D._600_ of 1 in 2 mL of LB with appropriate antibiotics. Cells were heat killed by incubating at 75°C on a heat block for 1 hour. Cells were then serially diluted in 1X TBS as described in the “alamarBlue cell viability assay” and “Colony forming units (CFU) assay” sections, and identical alamarBlue and CFU assays were performed to evaluate the metabolic and divisional capabilities of the heat killed cells. Cells were then stained with Rhodamine123 as described in the “Rhodamine123 staining” section, and the ratio of border-to-cytoplasm fluorescent signal was investigated with confocal microscopy as described in the “Rhodamine123 staining” section. Lastly, MAIM was performed on these cells as described in the “Multiple angle illumination microscopy (MAIM)” section, and the results were evaluated as described in the “MAIM image processing” section.

### Statistical analyses

Mann-Whitney t-tests were performed using GraphPad Prism version 10.3.1 for Windows, GraphPad Software, San Diego, California USA for all statistical analyses.

## Supporting information

All data are available in the manuscript or the supplementary materials

## Acknowledgments

The authors wish to thank the TU Delft 2016 iGEM team for the initial development and work done on this project: Lycka Kamoen, María Vázquez Vitali, Carmen Berends, Célina Reuvers, Charlotte Koster, Giannis Papazoglou, Iris de Vries, Lara van der Woude, Liza de Wilde, and Tessa Vergroesen; Matthew Hamilton Fyfe and Dr. Giulia Brachi from University of Colorado Boulder for laboratory support in Wil V. Srubar III’s laboratory; Morgan Brady from the University of Rochester for laboratory and project development and suggestions; Arjun Chaudhardy for troubleshooting on the heat-killing biological work; Karen Bentley, Chad Galloway, and Kelsea Cristillo of the Electron Microscopy Resource in the Center for Advanced Research Technologies at the University of Rochester Medical Center for preparing the cells for electron microscopy imaging; and the University of Rochester Medical Center’s Center for Advanced Light Microscopy and Nanoscopy, especially Julie Zhang and Kaye Thomas for their assistance with confocal microscopy.

## End notes

### Author Contributions

L.M.S. and A.S.M. conceived and planned the study. L.M.S. created the DNA constructs. L.M.S., K.P.S., M.M.B., D.F.W., and E.J. performed the microbiology characterization and microscopy analyses. E.A.A. developed the microscopy imaging techniques. B.C.A. and W.V.S. performed the SEM. All authors contributed to the writing of the final manuscript.

### Data and material availability

All data are available in the manuscript or the supplementary materials.

## References

1. Sidor, L. M. S., et al. Engineered bacterial that self-assemble bioglass polysilicate coatings display enhanced light focusing. Proc. Natl. Acad. Sci. U.S.A. 121 (51) e2409335121, (2024).

2. Aldea, M., Hernández-Chico, C., de la Campa, A. G., Kushner, S. R., and Vicente, M. Identification, cloning, and expression of bolA, an ftsZ-dependent morphogene of Escherichia coli. J. Bacteriol. 170, 5169–5176 (1988).

3. Dressaire, C., Moreira, R. N., Barahona, S., Alves de Matos, A. P., and Arraiano, C. M. BolA Is a Transcriptional Switch That Turns Off Motility and Turns On Biofilm Development. mBio 6, e02352–14 (2015).

4. Freire, P., Moreira, R. N., and Arraiano, C. M. BolA inhibits cell elongation and regulates MreB expression levels. J. Mol. Biol. 385, 1345–1351 (2009).

5. Santos, J. M., Freire, P., Vicente, M., and Arraiano, C. M. The stationary-phase morphogene bolA from Escherichia coli is induced by stress during early stages of growth. Mol. Microbiol. 32, 789–798 (1999).

6. Schoemaker, J. M., Gayda, R. C., and Markovitz, A. Regulation of cell division in Escherichia coli: SOS induction and cellular location of the sulA protein, a key to lon-associated filamentation and death. J. Bacteriol. 158, 551–561 (1984).

7. Vedyaykin, A., Rumyantseva, N., Khodorkovskii, M., and Vishnyakov, I. SulA is able to block cell division in *Escherichia coli* by a mechanism different from sequestration. Biochem. Biophys. Res. Commun. 525, 948–953 (2020).

8. Bean, G. J. et al. A22 Disrupts the Bacterial Actin Cytoskeleton by Directly Binding and Inducing a Low-Affinity State in MreB. Biochemistry 48, 4852–4857 (2009).

9. Iwai, N., Nagai, K., and Wachi, M. Novel *S*-Benzylisothiourea Compound That Induces Spherical Cells in *Escherichia coli* Probably by Acting on a Rod-shape-determining Protein(s) Other Than Penicillin-binding Protein 2. Biosci. Biotechnol. Biochem. 66, 2658–2662 (2002).

10. Altug, H., Oh, S.-H., Maier, S. A., and Homola, J. Advances and applications of nanophotonic biosensors. Nat. Nanotechnol. 17, 5–16 (2022).

11. Wu, F., van Schie, B. G. C., Keymer, J. E., and Dekker, C. Symmetry and scale orient Min protein patterns in shaped bacterial sculptures. Nat. Nanotechnol. 10, 719–726 (2015).

12. Pogliano, J., Pogliano, K., Weiss, D. S., Losick, R., and Beckwith, J. Inactivation of FtsI inhibits constriction of the FtsZ cytokinetic ring and delays the assembly of FtsZ rings at potential division sites. Proc. Natl. Acad. Sci. U. S. A. 94, 559–564 (1997).

13. Weiss, D. S., Chen, J. C., Ghigo, J.-M., Boyd, D., and Beckwith, J. Localization of FtsI (PBP3) to the Septal Ring Requires Its Membrane Anchor, the Z Ring, FtsA, FtsQ, and FtsL. J. Bacteriol. 181, 508–520 (1999).

14. Takeuchi, S., DiLuzio, W. R., Weibel, D. B.. and Whitesides, G. M. Controlling the Shape of Filamentous Cells of *Escherichia coli*. Nano Lett. 5, 1819–1823 (2005).

15. Chung, H. S. et al. Rapid β-lactam-induced lysis requires successful assembly of the cell division machinery. Proc. Natl. Acad. Sci. U. S. A. 106, 21872–21877 (2009).

16. Müller, W. et al. Bioencapsulation of living bacteria (*Escherichia coli*) with poly(silicate) after transformation with silicatein-α gene. Biomaterials 29, 771–779 (2008).

17. Li, C. W., Chu, S., and Lee, M. Characterizing the silica deposition vesicle of diatoms. IProtoplasmaI. 151, 158–163 (1989).

18. Nguyen, H. T. et al. Comparison of Two Transmission Electron Microscopy Methods to Visualize Drug-Induced Alterations of Gram-Negative Bacterial Morphology. Antibiotics 10, 307 (2021).

19. Rampersad, S. N. Multiple Applications of Alamar Blue as an Indicator of Metabolic Function and Cellular Health in Cell Viability Bioassays. Sensors 12, 12347–12360 (2012).

20. Darzynkiewicz, Z., Traganos, F., Staiano-Coico, L., Kapuscinski, J., and Melamed, M. R. Interaction of rhodamine 123 with living cells studied by flow cytometry. Cancer Res. 42, 799–806 (1982).

21. Mueller, E. A., Westfall, C. S., and Levin, P. A. pH-dependent activation of cytokinesis modulates Escherichia coli cell size. PLoS Genet. 16, e1008685 (2020).

22. Männik, J., Driessen, R., Galajda, P., Keymer, J. E., and Dekker, C. Bacterial growth and motility in sub-micron constrictions. Proc. Natl. Acad. Sci. U. S. A. 106, 14861–14866 (2009).

23. Rappé, M. S., Connon, S. A., Vergin, K. L., and Giovannoni, S. J. Cultivation of the ubiquitous SAR11 marine bacterioplankton clade. Nature 418, 630–633 (2002).

24. Schulz, H. N. et al. Dense populations of a giant sulfur bacterium in Namibian shelf sediments. Science 284, 493–495 (1999).

25. Volland, J.-M. et al. A centimeter-long bacterium with DNA contained in metabolically active, membrane-bound organelles. Science 376, 1453–1458 (2022).

26. Chavez, C. M. et al. The cell morphological diversity of Saccharomycotina yeasts. FEMS Yeast Res. 24, foad055 (2024).

27. Harris, J. M. et al. Long-Range Optogenetic Control of Axon Guidance Overcomes Developmental Boundaries and Defects. Dev. Cell 53, 577–588.e7 (2020).

28. Lee, S. et al. Inverse design of color routers in CMOS image sensors: toward minimizing interpixel crosstalk. Nanophotonics 13, 3895–3914.

29. Lorite, G. S. et al. Surface Physicochemical Properties at the Micro and Nano Length Scales: Role on Bacterial Adhesion and Xylella fastidiosa Biofilm Development. PLoS ONE 8, e75247 (2013).

30. Oh, S.-H. et al. Nanophotonic biosensors harnessing van der Waals materials. Nat. Commun. 12, 3824 (2021).

31. Li, Y. et al. Red-Blood-Cell Waveguide as a Living Biosensor and Micromotor. Adv. Funct. Mater. 29, 1905568 (2019).

32. Jones, D. & Elf, J. Bursting onto the scene? Exploring stochastic mRNA production in bacteria. Curr. Opin. Microbiol. 45, 124–130 (2018).

33. Part:BBa B0034 - parts.igem.org. https://parts.igem.org/wiki/index.php/Part:BBa_B0034.

34. Part:BBa K1890030 - parts.igem.org. https://parts.igem.org/wiki/index.php?title=Part:BBa_K1890030.

35. Guzman, L. M., Belin, D., Carson, M. J., and Beckwith, J. Tight regulation, modulation, and high-level expression by vectors containing the arabinose PBAD promoter. J. Bacteriol. 177, 4121–4130 (1995).

36. Part:pSB1C3 - parts.igem.org. https://parts.igem.org/Part:pSB1C3.

37. Schindelin, J., et al. Fiji: an open-source platform for biological-image analysis. Nat. Methods 9, 676–682 (2012).

