## Supplementary material for "Tunable light-focusing behavior of engineered bacterial microlenses with controllable shapes": All data are available in the manuscript or the supplementary materials

**This PDF file includes:**

Figs. S1 to S23
Supplemental Tables 1 and 2

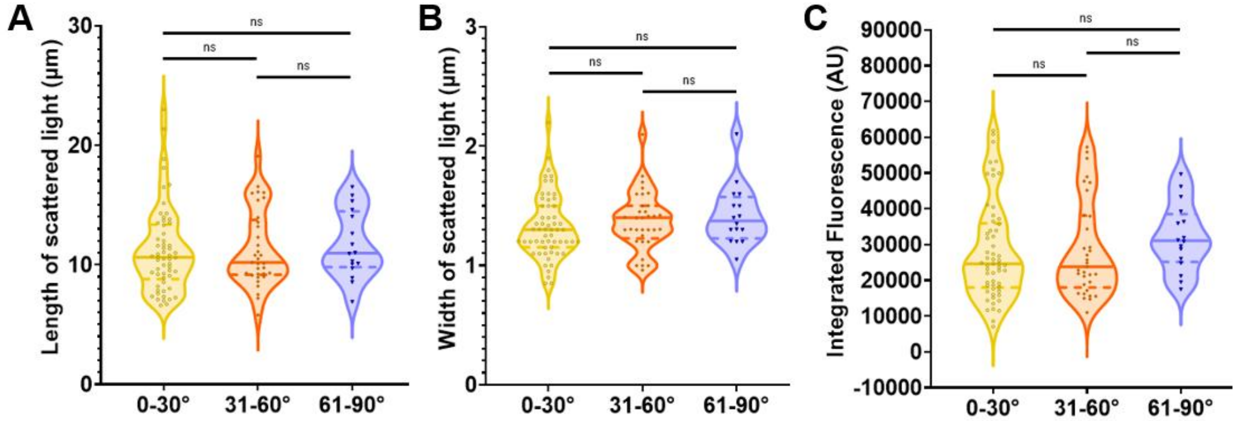


**Supplemental Figure 1: Scattered light length, width, and integrated intensity are not cell-orientation dependent for rod-shaped polysilicate-encapsulated bacteria.**
(A) Length of scattered light, (B) width of scattered light, and (C) integrated intensity of the scattered light, calculated from maximum intensity projections from Figure 1 for each angle bin. ($n_{0\text{-}30^\circ}$=58, $n_{31\text{-}60^\circ}$=33, $n_{61\text{-}90^\circ}$=16) ns: no significance

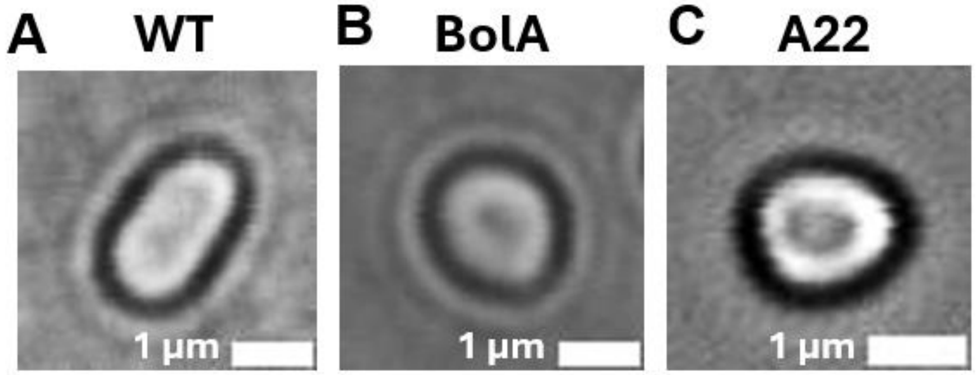


**Supplemental Figure 2: Shape-alteration of *E. coli* cells via BolA overexpression or A22 treatment.**
(A) Wild-type Top10 *E. coli* cell. (B) BolA-expressing *E. coli* cell. (C) A22-treated *E. coli* cell.

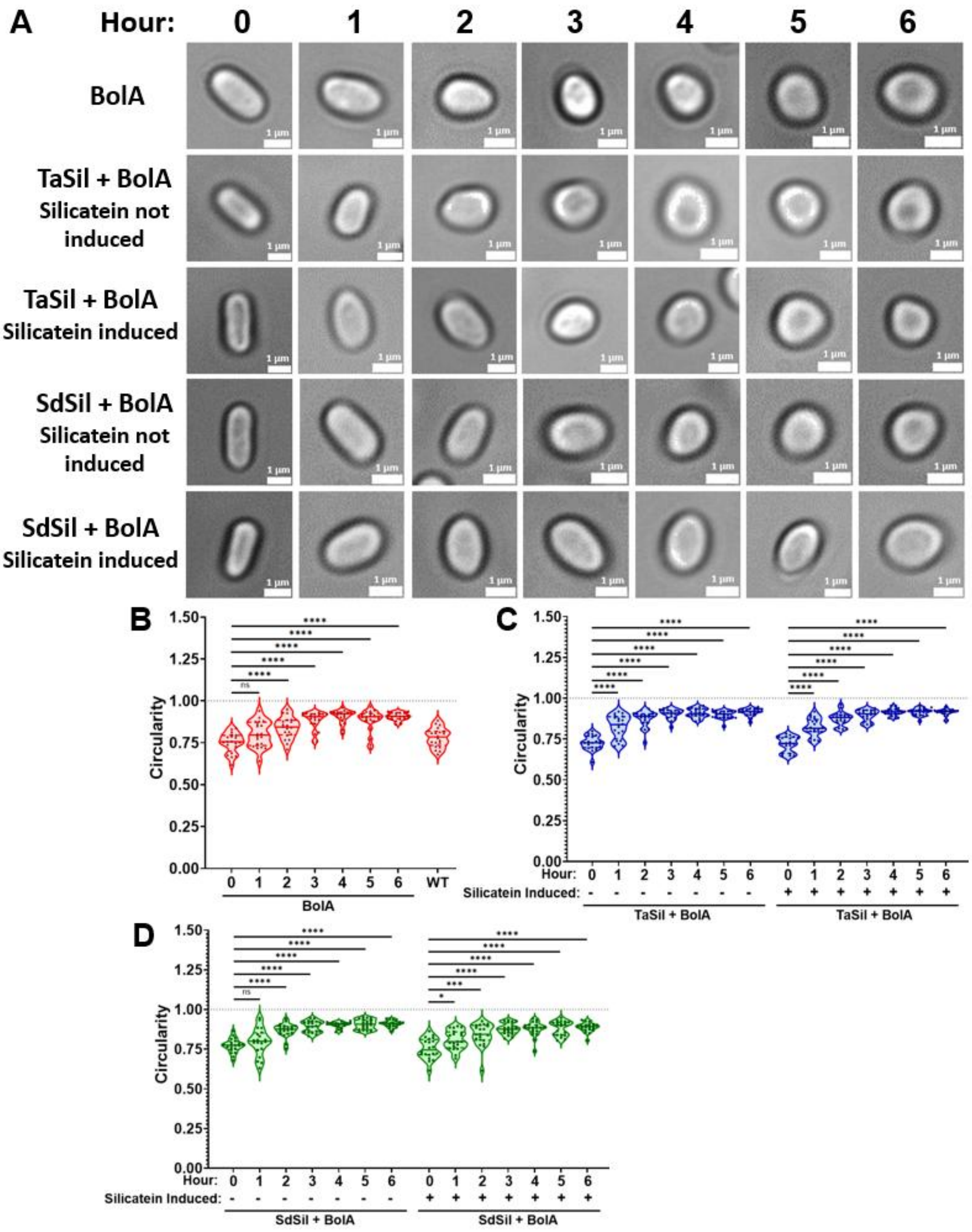


**Supplemental Figure 3: Time course of *E. coli* cell shape-change by BolA overexpression.** (A) Light microscopy images of BolA-expressing cells (BolA) following BolA induction, as well as BolA-expressing cells containing silicatein-expressing plasmids (TaSil+BolA and SdSil+BolA) following BolA induction and orthosilicate incubation, both with and without silicatein induction. (B-D) Quantification of the circularity of the (B) BolA, (C) TaSil+BolA, and (D) SdSil+BolA cells over time. (n=20) ns: no significance, * $p \leq 0.05$, *** $p \leq 0.001$, **** $p \leq 0.0001$.

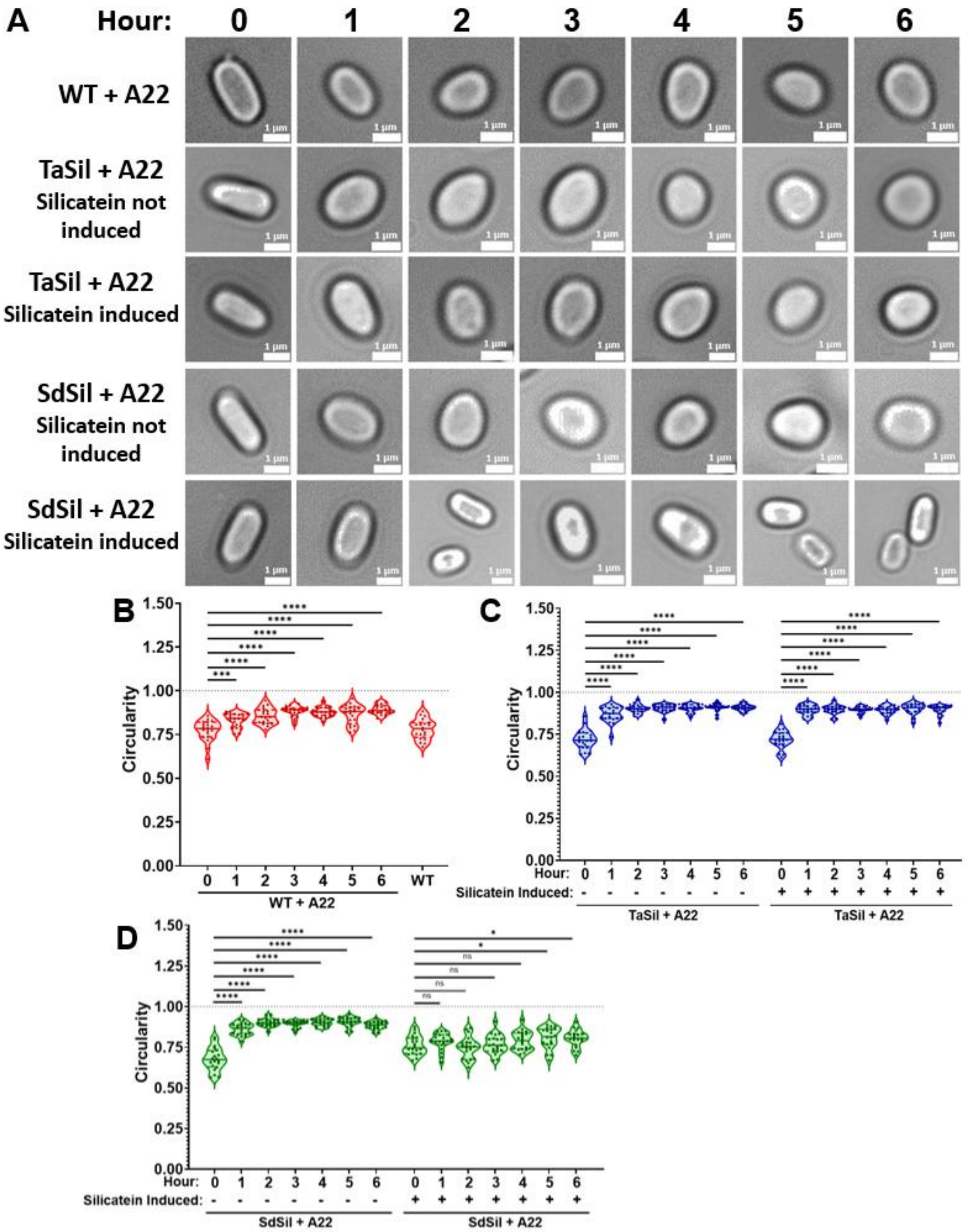


**Supplemental Figure 4: Time course of A22-treated *E. coli* cell shape-change.**
(A) Light microscopy images of A22-treated wild-type cells (WT+A22) as well as A22-treated silicatein-expressing cells (TaSil+A22 and SdSil+A22) following A22-treatment and orthosilicate incubation, both with and without silicatein induction. (B-D) Quantification of the circularity of the (B) WT+A22, (C) TaSil+A22, and (D) SdSil+A22 cells over time. (n=20) ns: no significance, * $p \leq 0.05$, *** $p \leq 0.001$, **** $p \leq 0.0001$.

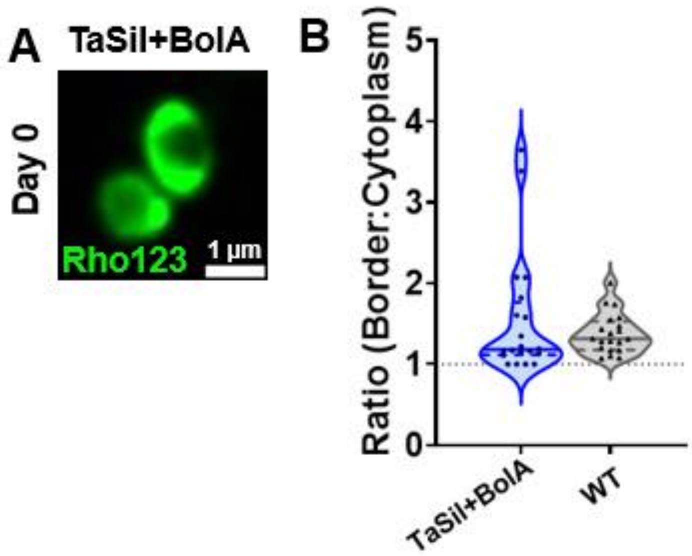


**Supplemental Figure 5: Polysilicate localization on spherical TaSil+BolA bacterial cells.**
(A) TaSil+BolA cells stained with Rhodamine123 at day 0. (B) The quantification of the ratio of Rhodamine123 border to cytoplasm staining for strains A and WT. (n=20)

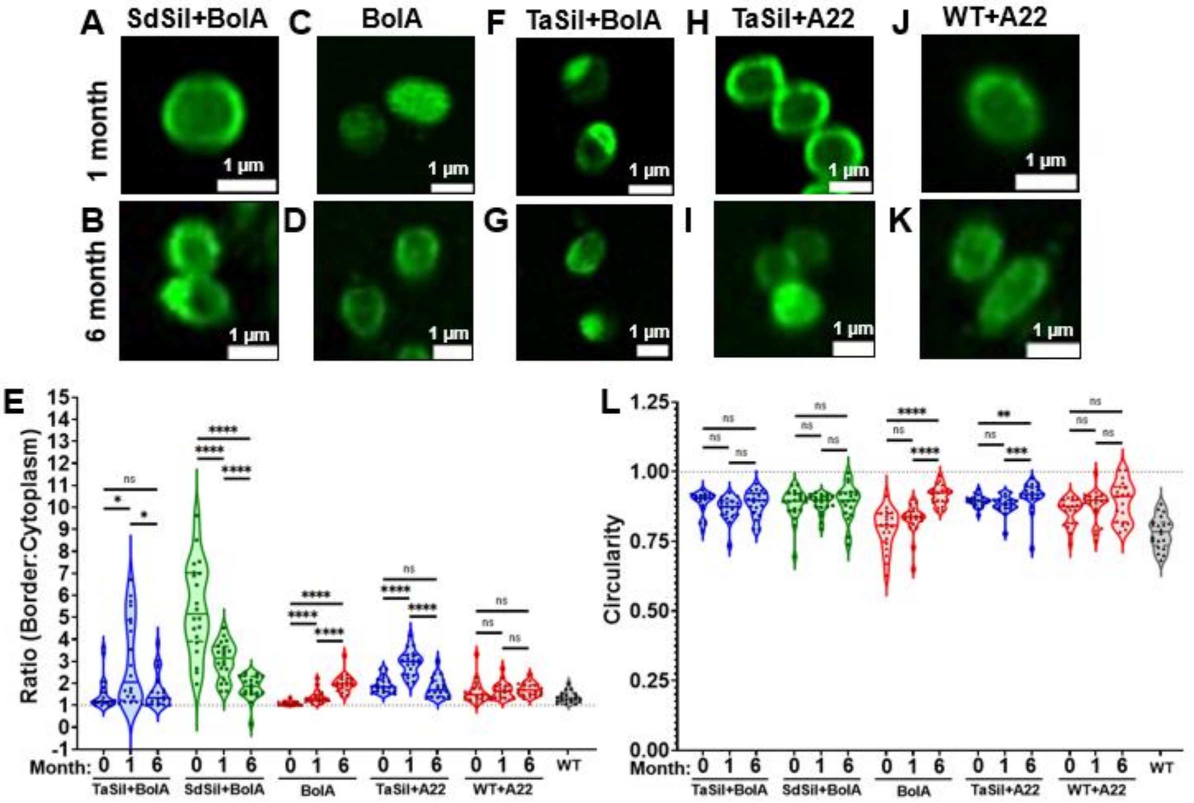


**Supplemental Figure 6: Polysilicate localization and circularity of spherical cells over time.** (A-D and F-K) Cells stained with Rhodamine123 at 1 month and 6 months where (A and B) are SdSil+BolA, (C and D) are BolA, (F and G) are TaSil+BolA, (H and I) is TaSil+A22, (J and K) is WT+A22. (E) The quantification of the ratio of Rhodamine123 border to cytoplasm staining for strains in Fig 2A-D and panels A-D and F-K. (L) Circularity quantifications for strains in Fig 2A-D and in panels A-D and F-K. (n=20) ns: no significance, * $p \leq 0.05$, ** $p \leq 0.01$, *** $p \leq 0.001$, **** $p \leq 0.0001$.

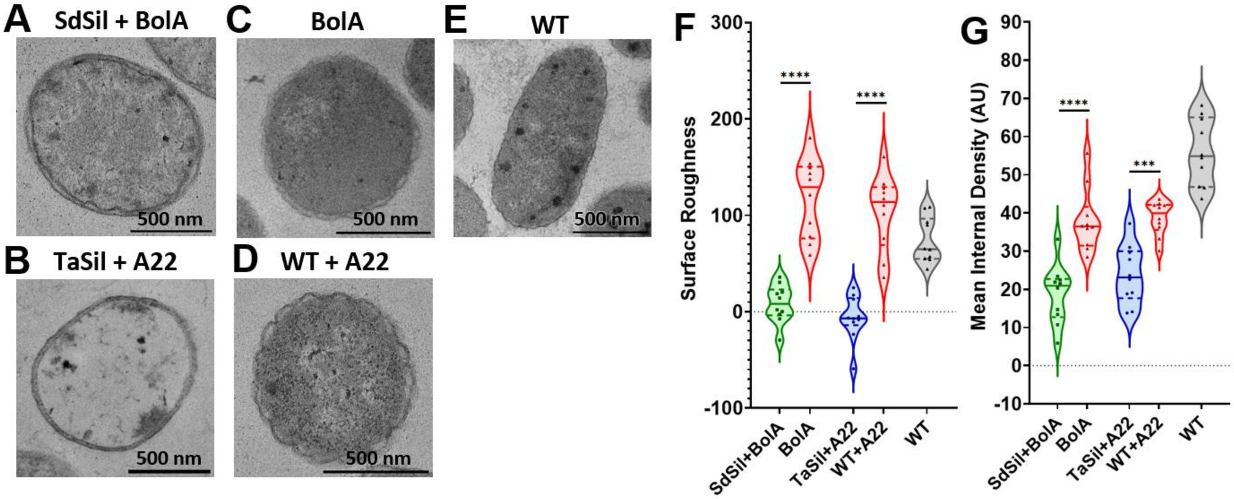


**Supplemental Figure 7: Polysilicate-encapsulated, spherical cells display smoother outer surfaces.**
(A-F) TEM images of thin-sections of the spherical, polysilicate-encapsulated cells, SdSil+BolA and TaSil+A22 (A and B), and non-encapsulated (C) BolA, (D) WT+A22, and (E) is WT. (F) Surface roughness quantification of cells in A-E. (G) Mean internal density quantification for cells in A-E. (n=10), *** $p \leq 0.001$, **** $p \leq 0.0001$.

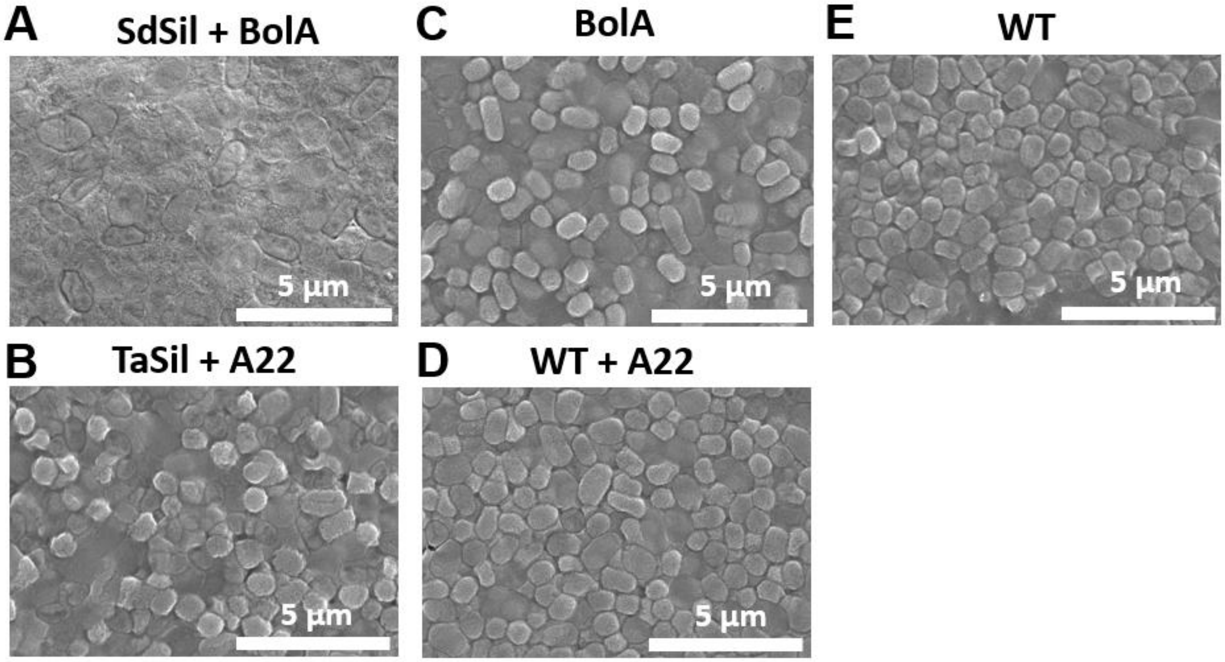


**Supplemental Figure 8: SEM of spherical shape-changed cells.**
(A-D) SEM images of (A) spherical, polysilicate-encapsulated SdSil+BolA *E. coli* cells, (B) spherical, polysilicate-encapsulated TaSil+A22 *E. coli* cells, (C) spherical BolA-expressing *E. coli* cells, (D) spherical WT+A22 *E. coli* cells, and (E) WT *E. coli* cells.

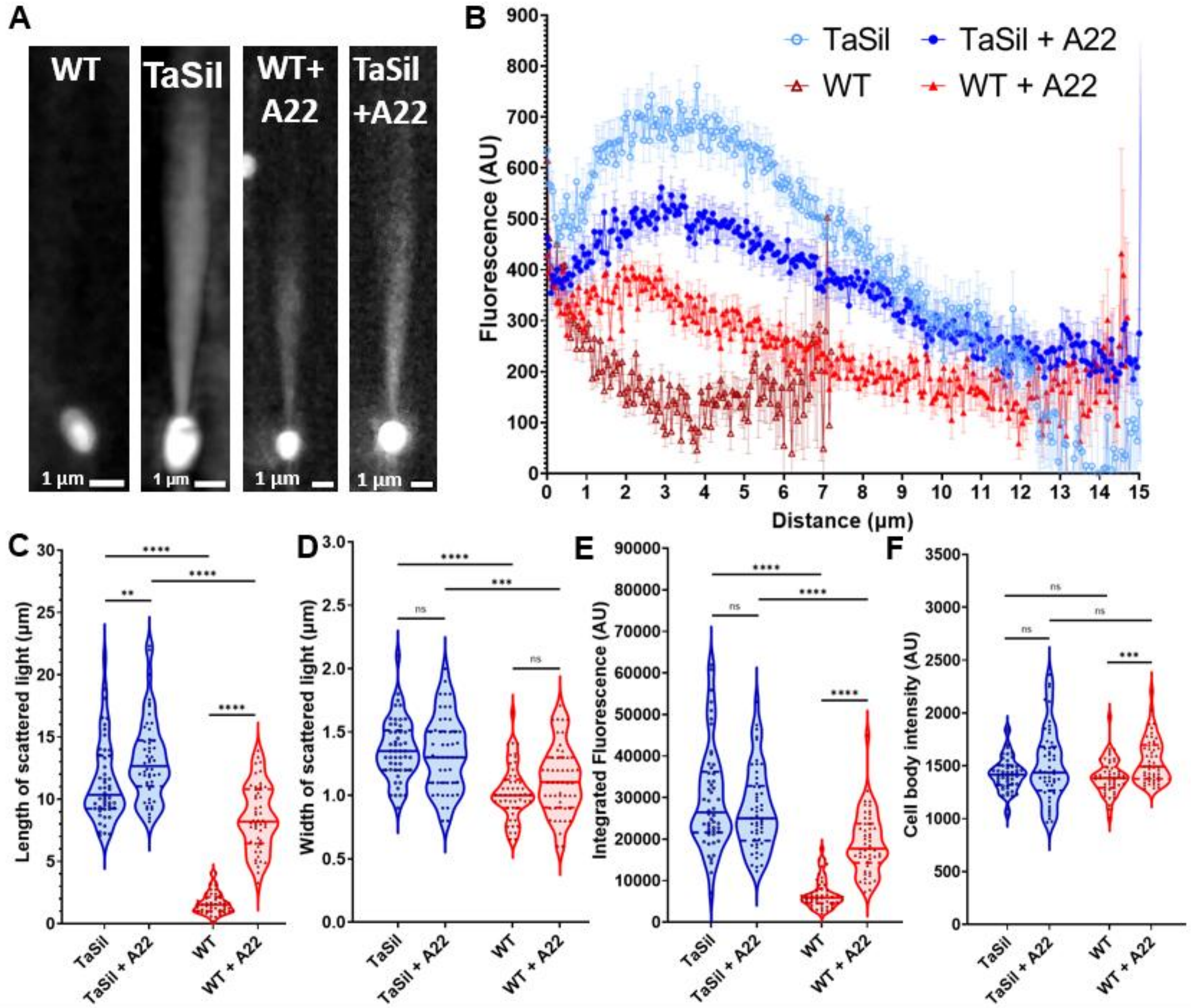


**Supplemental Figure 9: A22-treated spherical polysilicate-encapsulated cells scatter focused light that is less intense than rod-shaped polysilicate-encapsulated cells.**
(A) Maximum intensity projection for rod-shaped (WT), rod-shaped polysilicate-encapsulated (SdSil), spherical (WT+A22), and spherical polysilicate-encapsulated (TaSil+A22) cells scattering light via MAIM. (B) Profiles of the scattered light as a function of the distanced from the edge of the cell, calculated from the maximum intensity projections, where error bars correspond to standard error of the mean. (C) Length of the scattered light, (D) width of the scattered light, (E) integrated intensity of the scattered light, and (F) mean cell body intensity of the light within cell boundaries, all calculated from the maximum intensity projections. (n=50). ns: no significance, ** p≤0.01, *** p≤0.001, **** p≤0.0001.

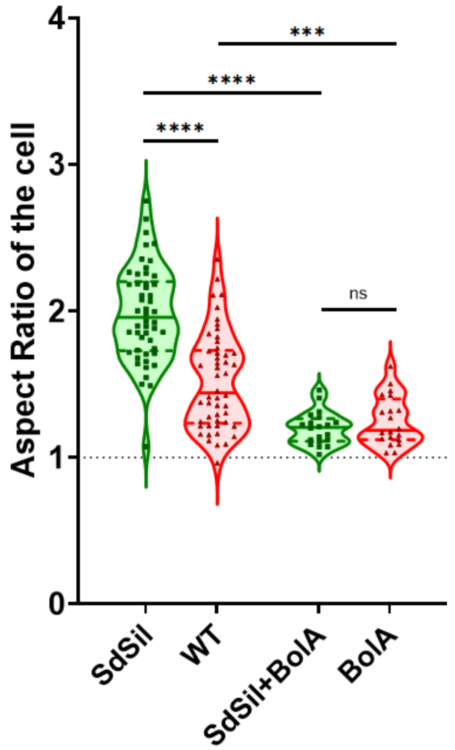


**Supplemental Figure 10: Aspect ratios of rod-shaped cells compared to spherical BolA-expressing cells.**

Aspect ratios calculated as length of the cell divided by the width of the cell for rod-shaped cells SdSil and WT, and round cells SdSil+BolA and BolA. Aspect ratio of 1 is circular, denotated by the dotted line. ($n_{\text{SdSil and WT}}$=50, $n_{\text{SdSil+BolA and BolA}}$=20) ns: no significance, *** $p \leq 0.001$, **** $p \leq 0.0001$

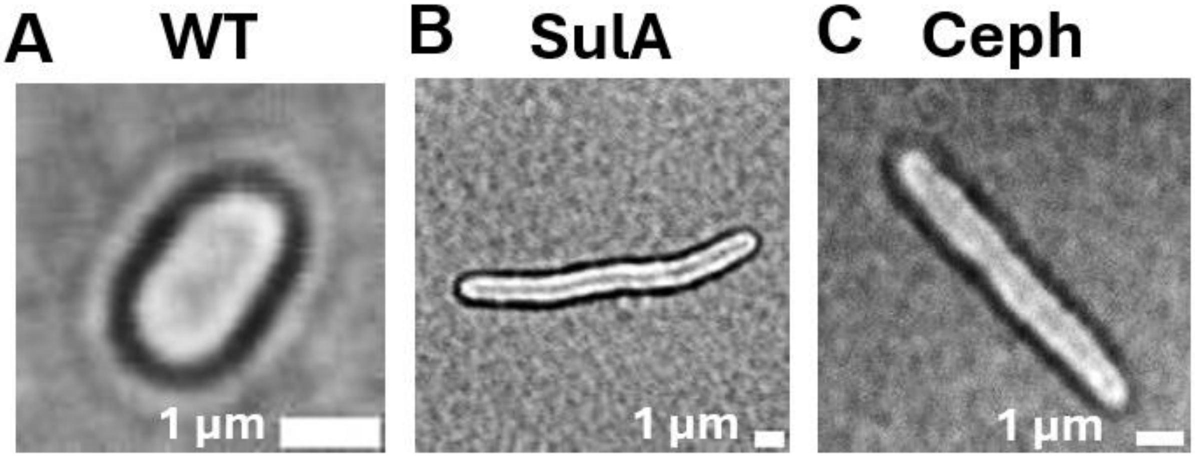


**Supplemental Figure 11: Shape alteration of *E. coli* cells via SulA overexpression or cephalexin treatment.**
(A) Wild-type Top10 *E. coli* cell. (B) SulA-expressing *E. coli* cell. (C) Cephalexin-treated *E. coli* cell.

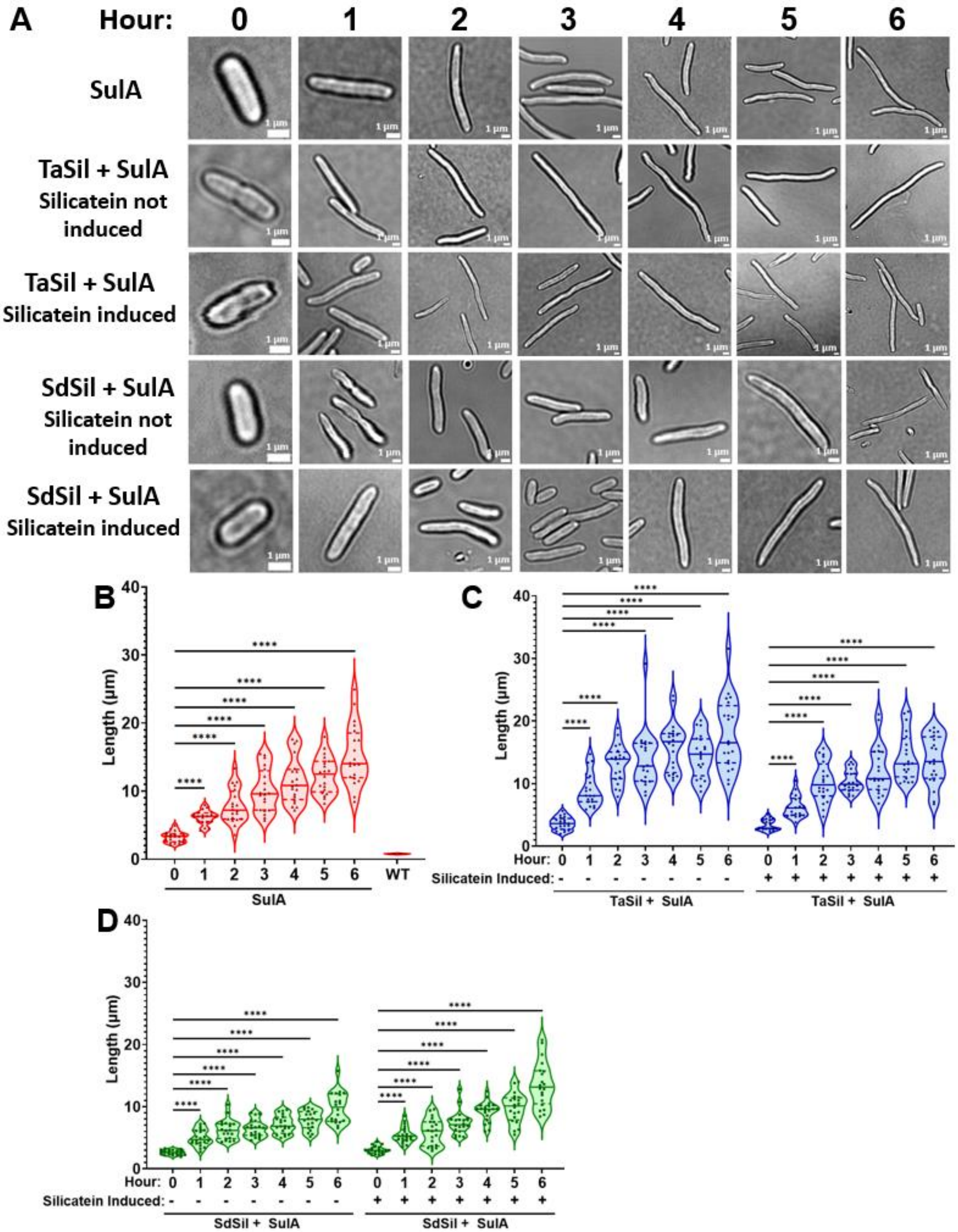


**Supplemental Figure 12: Time course of *E. coli* cell elongation by SulA overexpression.** (A) Light microscopy images of SulA-expressing cells (SulA) following SulA induction, as well as SulA-expressing cells containing silicatein-expressing plasmids (TaSil+SulA and SdSil+SulA) following SulA induction and orthosilicate incubation, both with and without silicatein induction. (B-D) Quantification of the circularity of the (B) SulA, (C) TaSil+SulA, and (D) SdSil+SulA cells over time. (n=20) **** p≤0.0001.

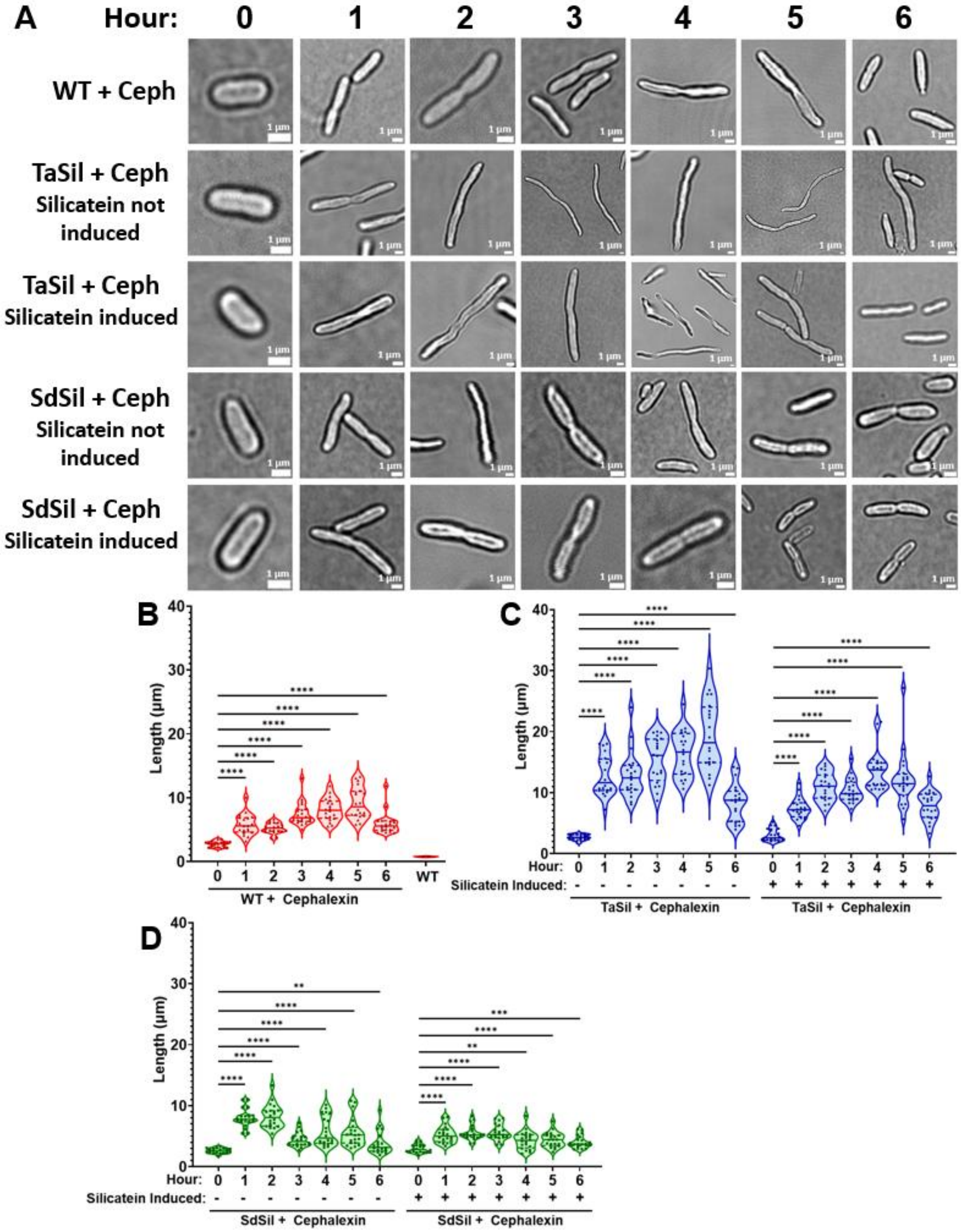


**Supplemental Figure 13: Time course of cephalexin-inducted *E. coli* cell elongation.**
(A) Light microscopy images of cephalexin-treated wild-type cells (WT+Ceph) as well as cephalexin-treated silicatein-expressing cells (TaSil+Ceph and SdSil+Ceph) following cephalexin-treatment and orthosilicate incubation, both with and without silicatein induction. (B-D) Quantification of the circularity of the (B) WT+Ceph, (C) TaSil+Ceph, and (D) SdSil+Ceph cells over time. (n=20) ** p≤0.01, *** p≤0.001, **** p≤0.0001.

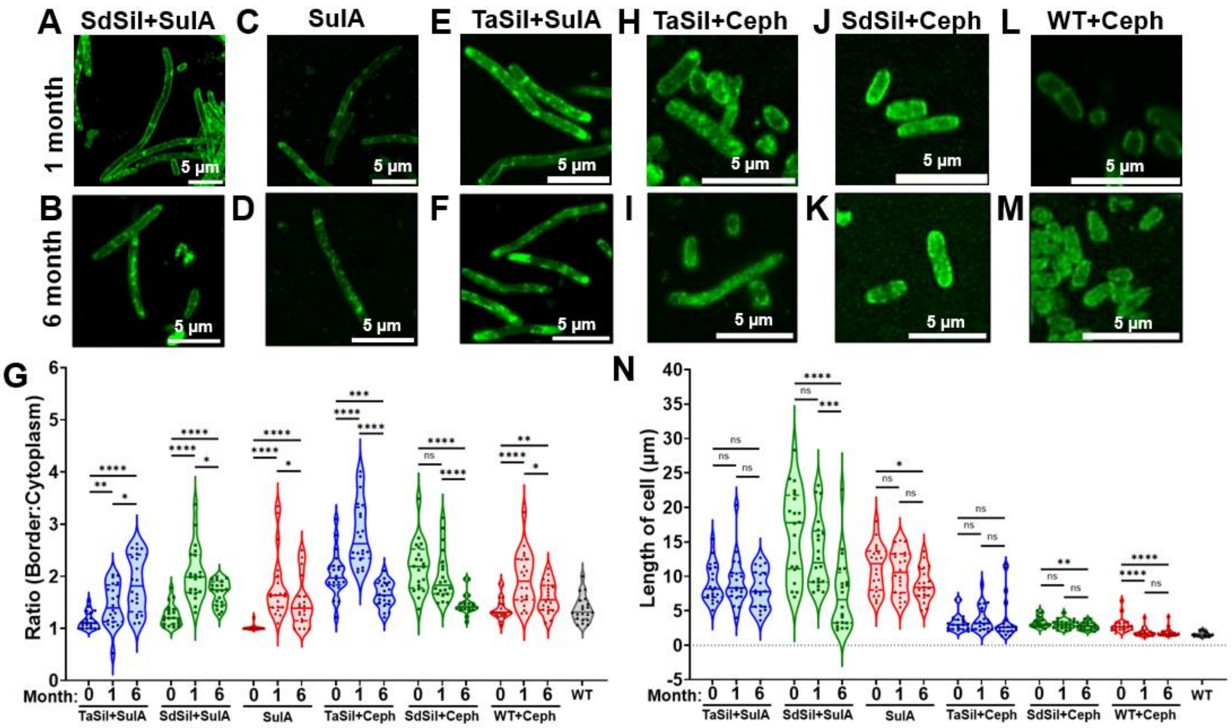


**Supplemental Figure 14: Polysilicate localization and length of elongated cells over time.** (A-F and H-M) Cells stained with Rhodamine123 at 1 month and 6 months where (A and B) are SdSil+SulA, (C and D) are SulA, (E and F) are TaSil+SulA, (H and I) are TaSil+Cephalexin, (J and K) are SdSil+Cephalexin, and (L and M) are WT+Cephalexin. (G) The quantification of the ratios of Rhodamine123 border to cytoplasm staining for strains in Fig 4A-C, Fig. 4E-G, and panels A-F and H-M. (N) Length quantifications for strains in Fig 4A-C, Fig. 4E-G and panels A-F and H-M. (n=20) ns: no significance, * $p\leq0.05$, ** $p\leq0.01$ *** $p\leq0.001$, **** $p\leq0.0001$.

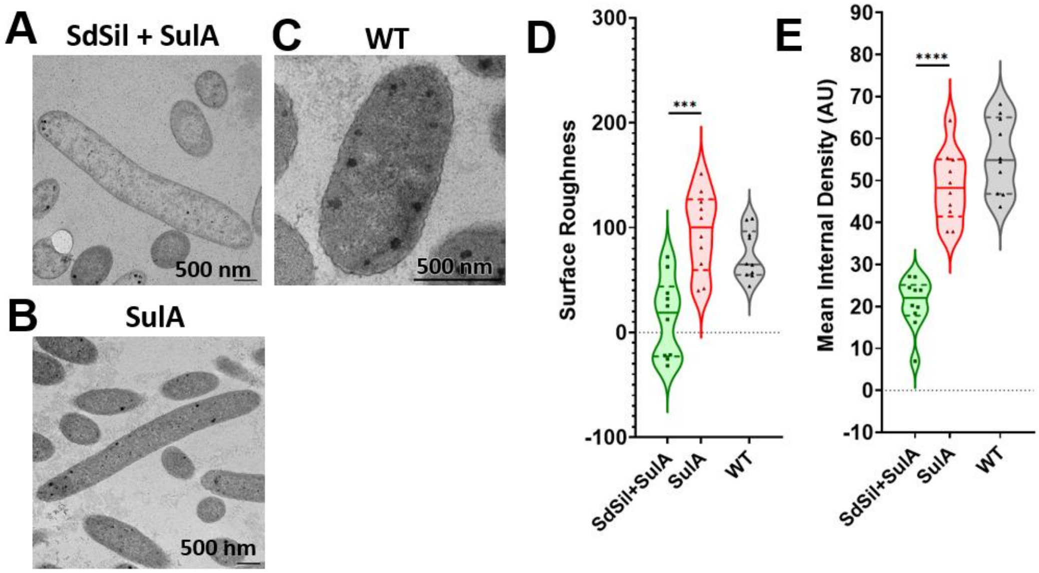


**Supplemental Figure 15: Polysilicate-encapsulated, elongated cells display smoother outer surfaces.**

(A-C) TEM images of thin-sections of (A) elongated, polysilicate-encapsulated SdSil+SulA cells, (B) non-encapsulated, elongated SulA cells, and (C) wild-type cells. (F) Surface roughness quantification of cells in A-C. (G) Mean internal density quantification of cells in A-C. (n=10), *** $p \leq 0.001$, **** $p \leq 0.0001$.

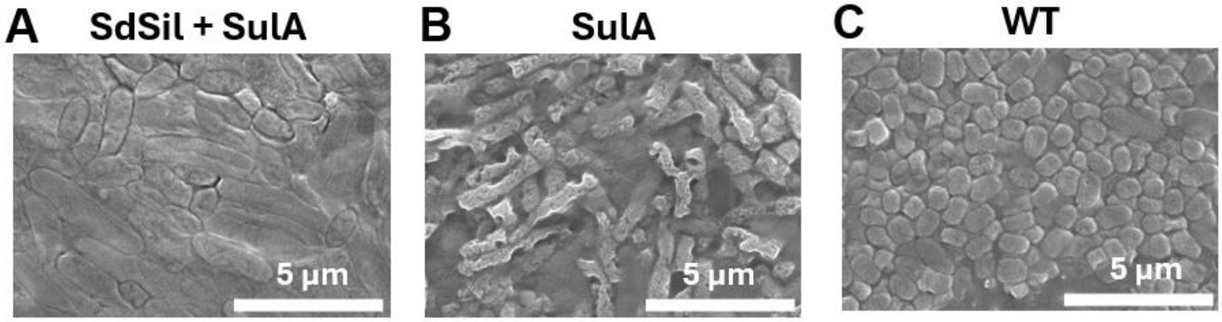


**Supplemental Figure 16: SEM of elongated, polysilicate-encapsulated cells.**
(A-D) SEM images of (A) highly-elongated, polysilicate-encapsulated SdSil+SulA, (B) highly elongated SulA, and (C) untreated wild-type.

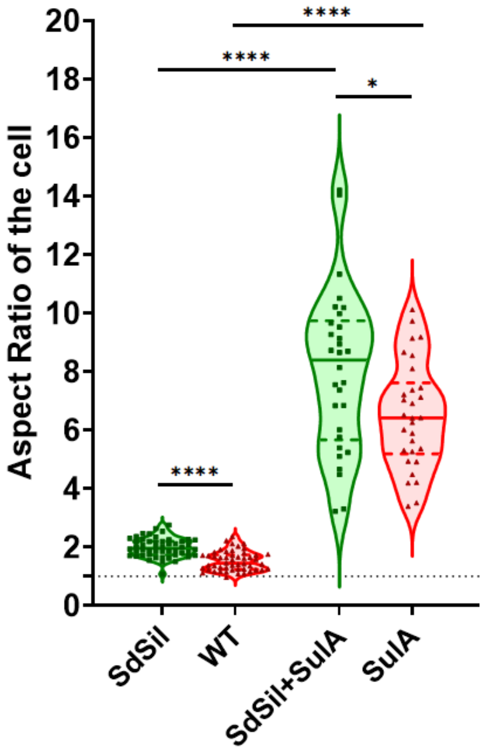


**Supplemental Figure 17: Aspect ratios of elongated cells compared to rod-shaped cells.** Aspect ratios calculated as length of the cell divided by the width of the cell for rod-shaped cells SdSil and WT, and highly elongated cells SdSil+SulA and SulA. Aspect ratio of 1 is circular, denotated by the dotted line. ($n_{\text{SdSil and WT}}$=50, $n_{\text{SdSil+SulA and SulA}}$=30) * $p \leq 0.05$, **** $p \leq 0.0001$

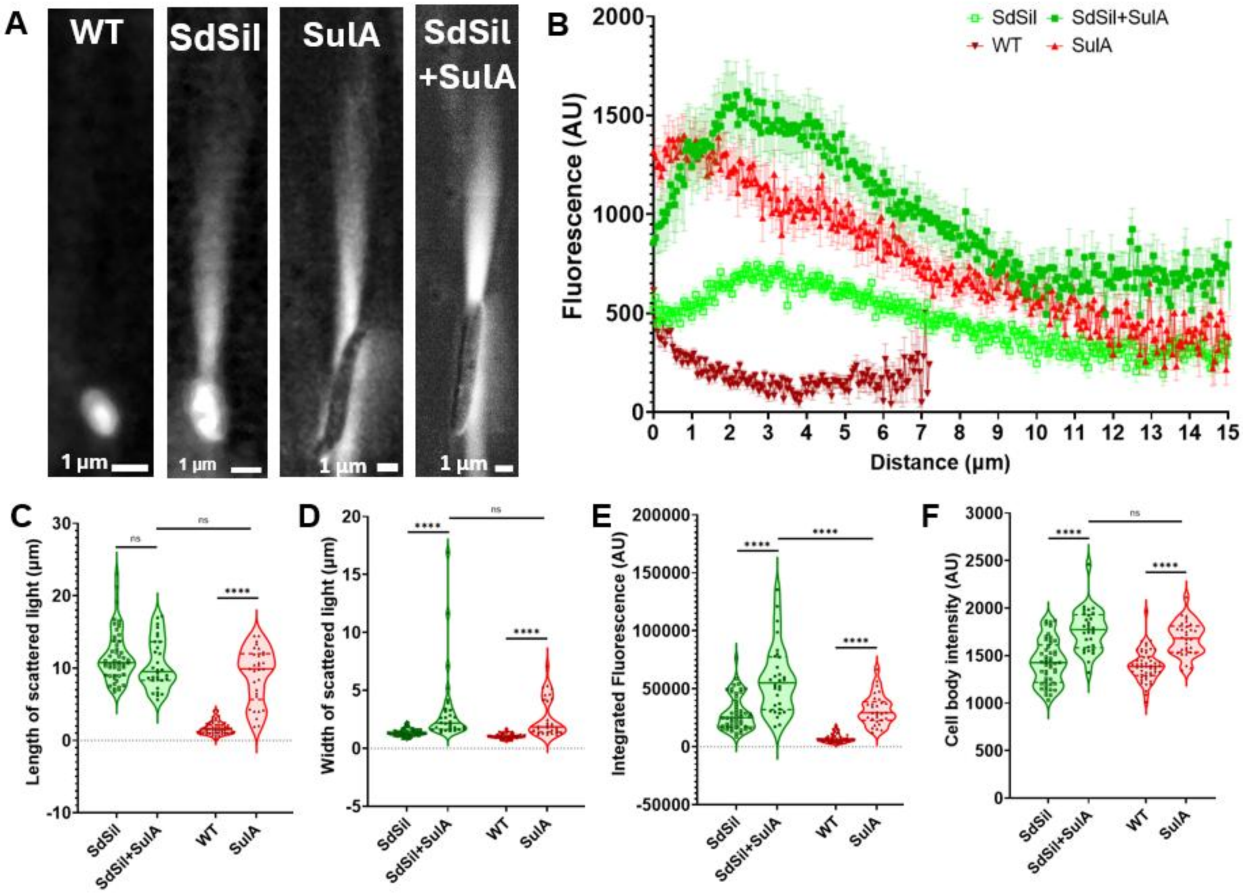


**Supplemental Figure 18: Elongated cells scatter more light than rod-shaped cells.**
(A) Maximum intensity projection for rod-shaped (WT), rod-shaped polysilicate-encapsulated (SdSil), highly elongated (SulA), and highly elongated, polysilicate-encapsulated (SdSil+SulA) cells scattering light via MAIM. (B) Profiles of the scattered light as a function of the distanced from the edge of the cell, calculated from the maximum intensity projections, where error bars correspond to standard error of the mean. (C) Length of the scattered light, (D) width of the scattered light, (E) integrated intensity of the scattered light, and (F) mean cell body intensity of the light within cell boundaries, all calculated from the maximum intensity projections. ($n_{\text{SdSil+SulA and SulA}}$=30, $n_{\text{SdSil and WT}}$ =50) ns: no significance, **** p≤0.0001.

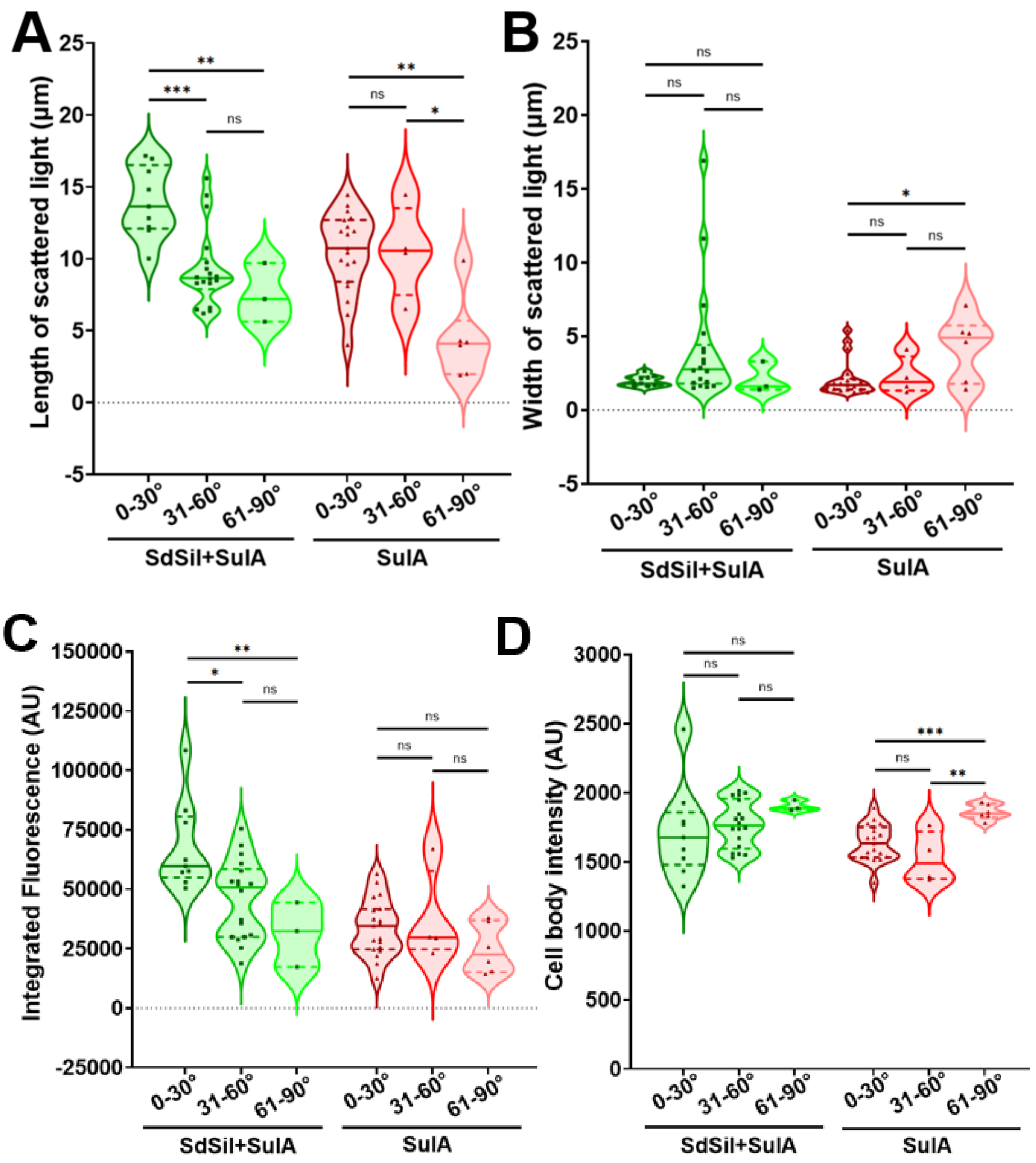


**Supplemental Figure 19: Quantification of the light scattering of elongated, polysilicate-encapsulated cells binned by angle relative to the incident light.**

(A) Length of the scattered light, (B) width of the scattered light, (C) integrated intensity of the scattered light, and (D) mean cell body intensity of the light within cell boundaries, all calculated from maximum intensity projections, with data binned by the angles of the cells relative to the incident light. ($n_{SulA,0\text{-}30°}$=19, $n_{SulA,31\text{-}60°}$=4, $n_{SulA,61\text{-}90°}$=6, $n_{SdSil+SulA,0\text{-}30°}$=9, $n_{SdSil+SulA,31\text{-}60°}$=18, $n_{SdSil+SulA,61\text{-}90°}$=3). ns: no significance, * $p \leq 0.05$, ** $p \leq 0.01$, *** $p \leq 0.001$.

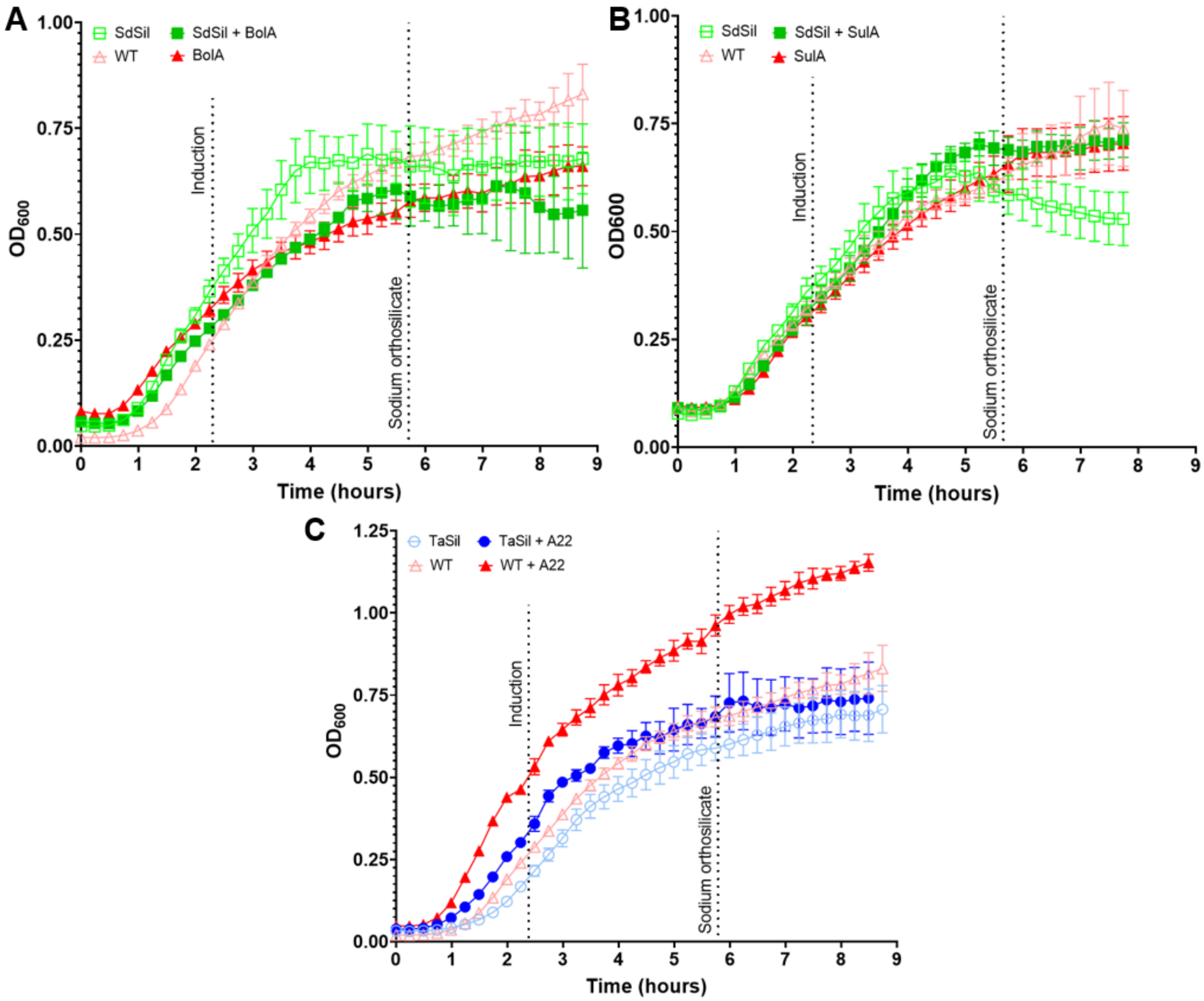


**Supplemental Figure 20: Growth curves for silicatein-expressing, shape-altered cells.** (A-C) Growth curves of silicatein-expressing, shape-altered strains over the time frame of the silicatein and shape-alteration induction and polysilicate encapsulation procedure. (A) Growth curves for wild-type (WT), silicatein-expressing (SdSil), spherical BolA-expressing (BolA), and spherical BolA- and silicatein-expressing (SdSil+BolA) strains. (B) Growth curves for wild-type (WT), silicatein-expressing (SdSil), elongated SulA-expressing (SulA), and elongated SulA- and silicatein-expressing (SdSil+SulA) strains. (C) Growth curves for wild-type (WT), silicatein-expressing (TaSil), spherical A22-treated (WT+A22), and spherical A22-treated and silicatein-expressing (TaSil+A22) strains.

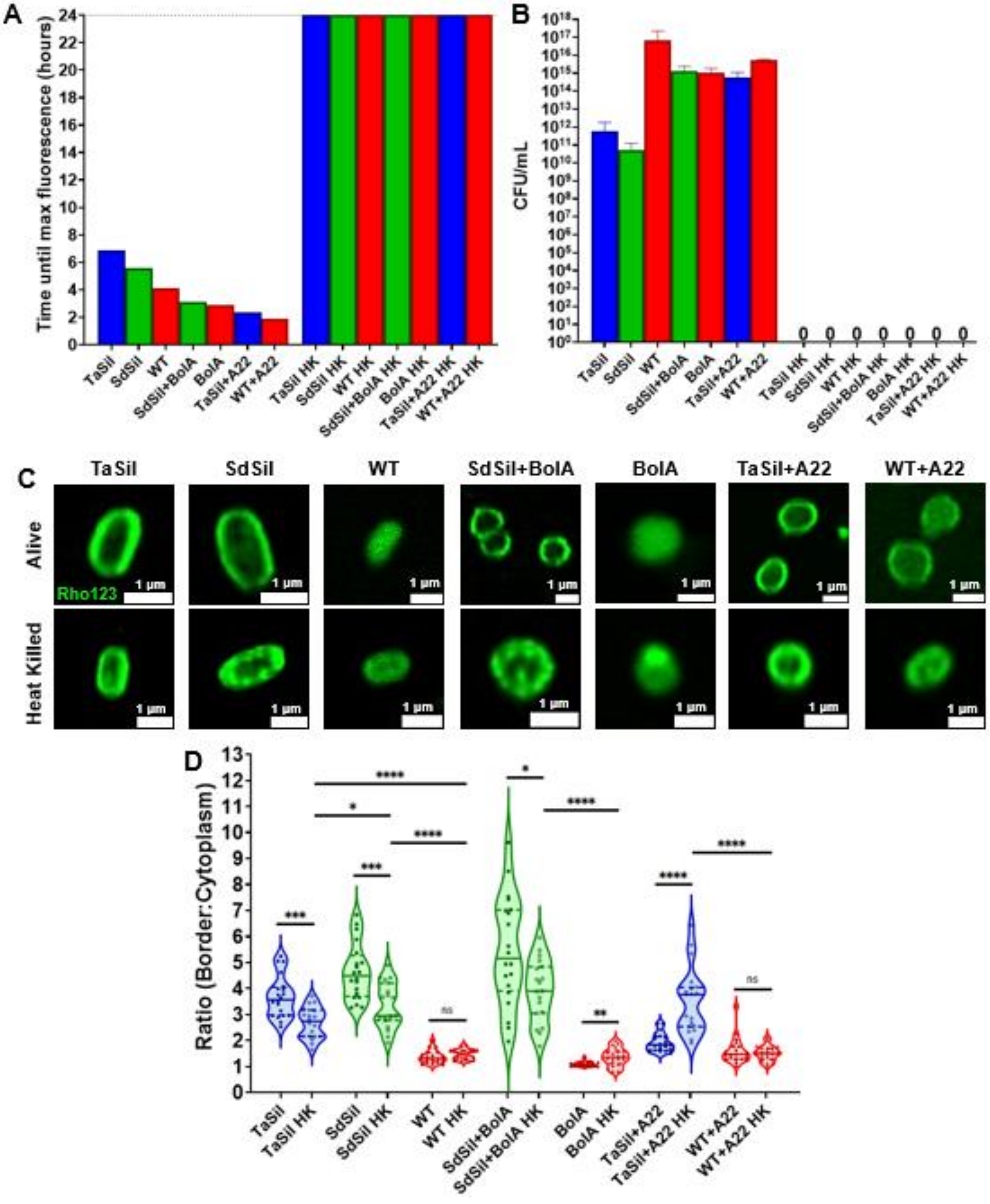


**Supplemental Figure 21: Heat-killed cells maintain polysilicate encapsulation morphology.** (A) Colony forming unit assays and (B) time until maximum fluorescence for alamarBlue metabolic assays for cells that were either freshly prepared or heat killed (HK) at 75°C for 1 hour. (C) Cells stained with Rhodamine123, where the top row cells are freshly prepared cells and the bottom row are cells that have been heat killed at 75°C for 1 hour. (D) Quantification of the ratios of Rhodamine123 border to cytoplasm staining for strains in C. ($n_{Alive}$=20, $n_{Heat\ Killed}$ = 20) ns: no significance, * $p \leq 0.05$, ** $p \leq 0.01$, *** $p \leq 0.001$, **** $p \leq 0.0001$.

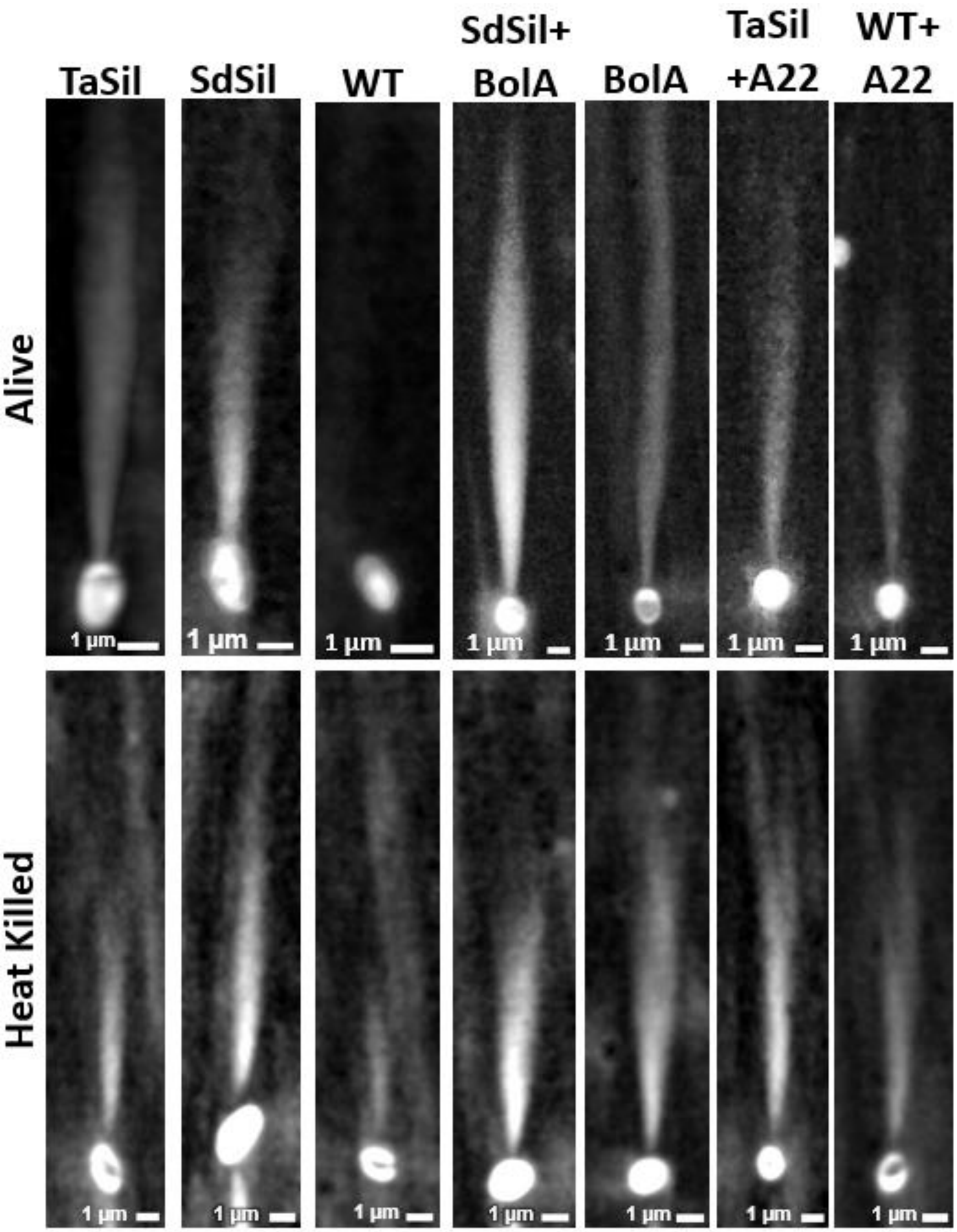


**Supplemental Figure 22: Light scattering via MAIM of live and heat-killed cells.** Maximum intensity projections for freshly prepared cells (top row) and heat-killed cells (bottom row) scattering light via MAIM.

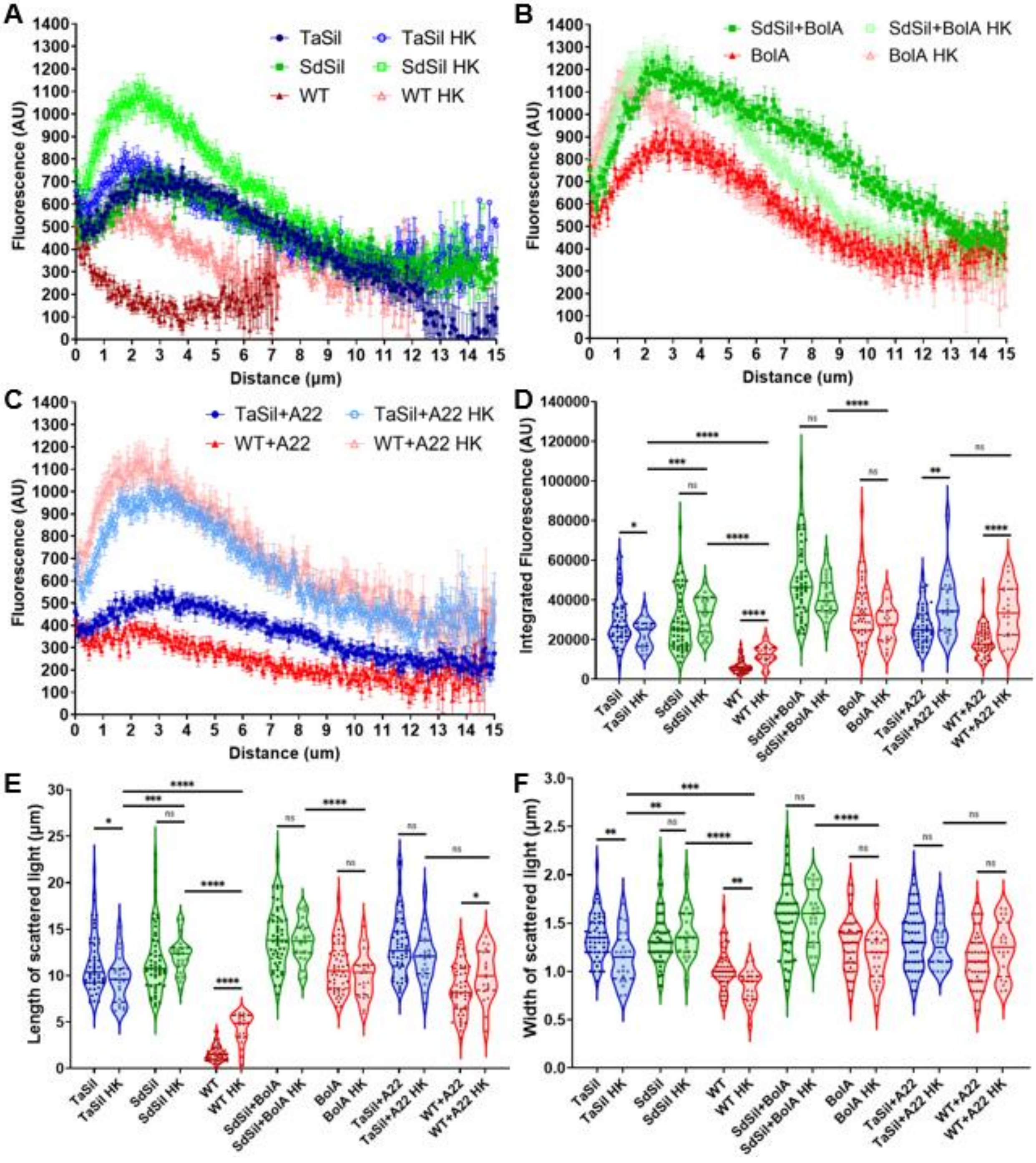


**Supplemental Figure 23: Heat killed cells scatter equivalent or more light than live cells.** (A, B, and C) Intensity of scattered light as a function of distance from the edge of the cell, calculated from maximum intensity projections via MAIM, for freshly prepared and heat-killed *E. coli* cells, where error bars correspond to the standard error of the mean. (D) Integrated intensity of the scattered light, (E) length of the scattered light, and (F) width of the scattered light, all calculated from the maximum intensity projections. ($n_{Alive}$=50, $n_{Heat\ Killed}$ = 20) ns: no significance, * p≤0.05, ** p≤0.01, *** p≤0.001, **** p≤0.0001.

**Supplemental Table 1: Shape-alteration approaches for silicatein-expressing strains**

| Silicatein Plasmid | Shape Change Method | Resulting Shape |
|---|---|---|
| TaSil | BolA pBAD33 | Spherical |
| | A22 | |
| | SulA pSB1C3 | Elongated |
| | Cephalexin | |
| SdSil | BolA pBAD33 | Spherical |
| | A22 | |
| | SulA pSB1C3 | Elongated |
| | Cephalexin | |

**Supplemental Table 2: Plasmid Information**

| Plasmid Name | Gene(s) of Interest | Vector | Resistance (final concentration) | Inducer (final concentration) |
|---|---|---|---|---|
| TaSil | OmpA-*Tethya aurantia* Silicatein | pBbS5a | Ampicillin (AMP) (100 µg/mL) | Isopropyl ß-D-1-thiogalactopyranoside (IPTG) (1 mM) |
| SdSil | OmpA-*Suberites domuncula* Silicatein | pRHA113 | AMP (100 µg/mL) | Rhamnose (0.2%) |
| BolA | BolA | pBAD33 | Chloramphenicol (CAM) (25 µg/mL) | Arabinose (30 mM) |
| SulA | SulA | pSB1C3 | CAM (25 µg/mL) | Arabinose (30 mM) |
